# Characterization of the hepatitis E virus replicase

**DOI:** 10.1101/2021.12.03.471094

**Authors:** Karoline Metzger, Cyrine Bentaleb, Kévin Hervouet, Virginie Alexandre, Claire Montpellier, Jean-Michel Saliou, Martin Ferrié, Charline Camuzet, Yves Rouillé, Cécile Lecoeur, Jean Dubuisson, Laurence Cocquerel, Cécile-Marie Aliouat-Denis

**Affiliations:** Univ. Lille, CNRS, INSERM, CHU Lille, Institut Pasteur de Lille, U1019 - UMR 9017 – CIIL - Center for Infection and Immunity of Lille, F-59000 Lille, France; Univ. Lille, CNRS, Inserm, CHU Lille, Institut Pasteur de Lille, UMR2014 - US41 - PLBS-Plateformes Lilloises de Biologie & Santé, Lille, France

**Keywords:** HEV p6, positive-strand RNA virus, ORF1 processing, *Gaussia* luciferase replicon, epitope tag, RNA hybridization, endocytic recycling compartment, replication complexes

## Abstract

Hepatitis E virus (HEV) is the major cause of acute hepatitis worldwide. HEV is a positive-sense RNA virus expressing 3 open reading frames (ORFs). ORF1 encodes the ORF1 non– structural polyprotein, the viral replicase which transcribes the full-length genome and a subgenomic RNA that encodes the structural ORF2 and ORF3 proteins. The present study is focused on the replication step with the aim to determine whether the ORF1 polyprotein is processed during the HEV lifecycle and to identify where the replication takes place inside the host cell. As no commercial antibody recognizes ORF1 in HEV-replicating cells, we aimed at inserting epitope tags within the ORF1 protein without impacting the virus replication efficacy. Two insertion sites located in the hypervariable region were thus selected to tolerate the V5 epitope while preserving HEV replication efficacy. Once integrated into the infectious full-length Kernow C-1 p6 strain, the V5 epitopes did neither impact the replication of genomic nor the production of subgenomic RNA. Also, the V5-tagged viral particles remained as infectious as the wildtype particles to Huh-7.5 cells. Next, the expression pattern of the V5-tagged ORF1 was compared in heterologous expression and replicative HEV systems. A high molecular weight protein (180 kDa) that was expressed in all 3 systems and that likely corresponds to the unprocessed form of ORF1 was detected up to 25 days after electroporation in the p6 cell culture system. Additionally, less abundant products of lower molecular weights were detected in both in cytoplasmic and nuclear compartments. Concurrently, the V5-tagged ORF1 was localized by confocal microscopy inside the cell nucleus but also as compact perinuclear substructures in which ORF2 and ORF3 proteins were detected. Importantly, using *in situ* hybridization (RNAScope®), positive and negative-strand HEV RNAs were localized in the perinuclear substructures of HEV-producing cells. Finally, by simultaneous detection of HEV genomic RNAs and viral proteins in these substructures, we identified candidate HEV factories.

## Introduction

Hepatitis E virus (HEV) is one of the leading causes of acute hepatitis worldwide (WHO, 2021). Amongst the 20 million infections estimated by WHO every year, 3.3 million cases are symptomatic. Although HEV infection is usually self-resolving in the general population with a mortality rate of 0.5 to 4% due to fulminant hepatitis, the immunocompromised patients, mainly organ transplant recipients, may suffer from chronic hepatitis and cirrhosis (Lhomme et al., 2020). Elevated mortality rates (up to 25%) have also been recorded among pregnant women in developing countries as well as in patients with pre-existing liver diseases (Pérez Gracia et al., 2017; Lhomme et al., 2020; Webb and Dalton, 2020). In addition, both chronic and acute HEV infections can lead to neurological disorders or kidney injuries and impaired renal function (Lhomme et al., 2020; Webb and Dalton, 2020).

HEV is classified in the *Hepeviridae* family and the 4 genotypes (gt 1-gt 4) that account for most of the human infections, are included within the *Orthohepevirus A* genus (Smith and Simmonds, 2018). HEV gt 1 and 2 are transmitted from human-to-human through fecal-oral route and can cause large, primarily waterborne outbreaks in resource-limited settings. Ingestion of undercooked swine or game meat is the primary mode of zoonotic transmission of HEV gt 3 and 4 in middle- and high-income areas (Kamar et al., 2017).

The HEV genome is a positive-sense, 5’-capped, single-stranded RNA of ∼7.2 kb in length. It is organized into 3 open reading frames (ORFs): ORF1, ORF2, and ORF3 (Wang and Meng, 2021). ORF1 encodes the ORF1 nonstructural polyprotein, which contains several functional domains essential for viral replication. ORF2 encodes the ORF2 viral capsid protein, which is involved in particle assembly, binding to host cells and eliciting neutralizing antibodies (Schofield et al., 2000; Shiota et al., 2013). ORF3 encodes a small multifunctional phosphoprotein involved in virion morphogenesis and egress (reviewed in (Glitscher and Hildt, 2021). ORF2 and ORF3 are partially overlapping and the corresponding proteins are translated from a subgenomic RNA of 2.2 kb in length (Graff et al., 2006).

ORF1 is the largest ORF in the viral genome and encodes a non-structural polyprotein where several domains have been bioinformatically assigned based on homology with Rubi-like viruses, *i.e*. *Rubivirus*, *Betateravirus* and *Benyvirus* (Koonin et al., 1992). Although several domains such as methyltransferase (Met), helicase (Hel) and RNA-dependent RNA polymerase (RdRp) have been reported to be enzymatically active, the function of the Y and X-domains as well as the highly disordered hypervariable region (HVR) remain elusive (Wang and Meng, 2021) (**Figure 1C**). In addition, the precise location of the protease region (known as papain-like cysteine protease, PCP) and its enzymatic activity are still a matter of debate (LeDesma et al., 2019; Proudfoot et al., 2019). Whether or not the HEV polyprotein gets processed by the PCP or cellular proteases remains a difficult question to address considering the low expression level of the polyprotein in HEV cell culture systems as well as the scarcity of functional specific antibodies (Debing et al., 2016; Lenggenhager et al., 2017; Nimgaonkar et al., 2018; Kenney and Meng, 2019).

**Figure 1.**
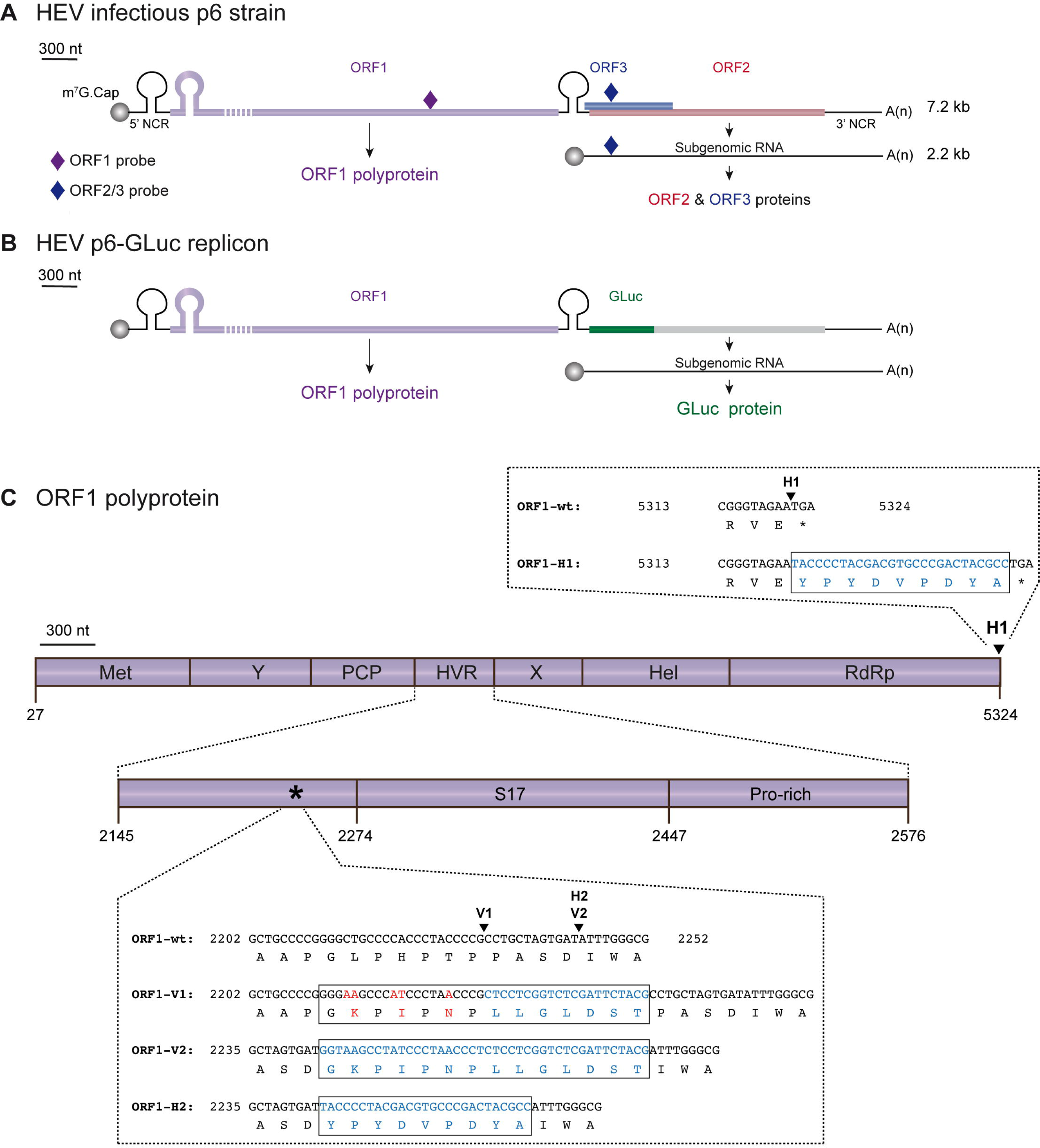
Localization scheme of epitope tag insertions within the ORF1 polyprotein of HEV. **Figure 1A.** Scheme of the HEV Kernow C-1 p6 (GenBank accession number JQ679013) genome and encoded proteins. The full-length HEV genome (7.2 kb) is composed of 5’ and 3’ non coding regions (NCR) as well as 3 open reading frames: ORF1 (purple) encoding the HEV replicase, ORF2 (red) and ORF3 (blue) encoding the capsid and a small phosphoprotein, respectively. The HEV subgenomic RNA of 2.2kb is depicted as a thin black line. The probes used in RT-qPCR experiments are located as small colored rhombus: ORF1-specific probe (dark purple) and ORF2/ORF3-specific probe (dark blue). The full-length of the ORF1 protein cannot be represented at the scheme scale (purple dashed lines). **Figure 1B.** Scheme of the HEV p6-GLuc replicon genome and encoded proteins. The *Gaussia* luciferase (GLuc, green) replaces the ORF3 and the N-terminus of ORF2 in the HEV p6-GLuc replicon. **Figure 1C.** The ORF1 is enlarged and its functional domains are designated as methyltransferase (MeT), Y domain (Y), papain-like cysteine protease (PCP), hypervariable region (HVR), macro- or X-domain (X), helicase (Hel), RNA-dependent RNA polymerase (RdRp). Insertion position of an HA epitope at the C-terminus of ORF1 is localized (H1, arrowhead). C-terminal nucleotide and aa sequences of RdRp are detailed for the ORF1 untagged wildtype (ORF1-wt) and HA-tagged (ORF1-H1) proteins with inserted epitope sequence highlighted in blue (box). A focus on the HVR enables to locate the human S17 ribosomal protein insertion (S17) of the HEV Kernow C-1 p6 strain and the prolin-rich region (Pro-rich). An asterisk points at the 2202-2252 region whose nucleotide and aa sequences are detailed (box). Two other sites of insertion of V5 or HA epitopes are shown (arrowheads): V1 and V2 for V5 epitope insertions and H2 for HA epitope insertion. Part of the HVR sequences of the wild-type (ORF1-wt) as well as the epitope-tagged ORF1 are presented (ORF1-V1, ORF1-V2, ORF1-H2). Mutated nucleotides and aa residues are in red. Inserted nucleotides and aa residues are in blue.

In this study, we sought to characterize the ORF1 replicase. The insertion of epitope tags into the HEV replicase came to us as the best strategy to track the polyprotein inside the host cell and identify its potential cleavage products. The HVR was chosen based on amino acid (aa) ORF1 sequence alignments as well as its capacity to tolerate inserted fragments arising either from duplication of viral genome or from human genes (Shukla et al., 2011, 2012; Nguyen et al., 2012; Johne et al., 2014; Lhomme et al., 2014). Additionally, the HVR region of the HEV83-2-27 strain of gt 3 has been recently reported to tolerate the insertion of an HA epitope and a small luciferase reporter gene (Szkolnicka et al., 2019). In our study, we attempted to insert V5 or HA epitopes into the HVR of gt 3 Kernow C-1 p6 strain, taking advantage of the homology existing between the HEV and V5 epitope aa sequences and also of a naturally occurring insertion in the HEV genome (Nguyen et al., 2012) (**Figure 1C**). At first, a p6-GLuc replicon expressing the *Gaussia Luciferase* (*GLuc*) reporter gene under the control of the epitope tagged-ORF1 was used to select insertion sites that did not impact replication efficacy (Shukla et al., 2012; Emerson et al., 2013) (**Figure 1B**). Two positions located into the HVR were selected as the V5 insertions did neither impact the *Gaussia* luciferase secretion nor the transcription of genomic and subgenomic viral RNAs. Also, the V5-tagged viral particles remained as infectious as the wildtype particles to Huh-7.5 cells. Next, the expression of the selected epitope-tagged ORF1 was analyzed in the p6-GLuc replicon, heterologous and p6 cell culture systems. The full-length ORF1 protein as well as less abundant products of lower molecular weight were detected in all systems, both in cytoplasmic and nuclear compartments. Simultaneous detection of HEV genomic / subgenomic RNAs by fluorescence *in situ* hybridization (RNAscope®) and ORF1 protein by immunofluorescence identified candidate HEV replication complexes as compact perinuclear nuggets in which ORF2 and ORF3 proteins were also detected. Finally, partial co-localization of viral proteins with cellular markers of the endocytic recycling compartment (ERC) unveiled the composition of the HEV replication factories.

## Materials and methods

### Plasmids

The plasmid pBlueScript SK (+) carrying the DNA of the full-length genome of HEV gt 3 Kernow C-1 p6 strain (GenBank accession number JQ679013) was kindly provided by Dr S.U. Emerson (**Figure 1A**). The HEV p6-wt-GLuc replicon, constructed from the HEV gt 3 Kernow C-1 p6 strain was also obtained from Dr Emerson (**Figure 1B**). This replicon possesses a *Gaussia Luciferase* (*GLuc*) reporter gene that substitutes the 5’ part of the ORF2 gene and most part of the ORF3 gene (Shukla et al., 2012; Emerson et al., 2013). A p6-GAD-GLuc mutant replicon in which the ORF1 polymerase active site GDD was mutated to GAD to prevent any replication was used as a negative control (Emerson et al., 2013).

The plasmids pTM-ORF1, pTM-ORF3/2 and pTM empty vector were kindly provided by Dr J. Gouttenoire from the University of Lausanne (Switzerland) and have been previously described (Lenggenhager et al., 2017). The pTM-ORF1 vector contains the full-length sequence of the ORF1 protein. The pTM empty vector, was used as a negative control.

### Epitope tag insertions

The HA and V5 epitopes were inserted in 3 different positions of the ORF1 sequence (**Figure 1C**). According to the insertion position, the constructs harboring the HA epitope are named H1 or H2 and the constructs harboring the V5 epitope are named V1 or V2. The pBlueScript SK(+) harboring the p6-GLuc replicon was used as a template. Sequences of the primers used to make the tagged ORF1 constructs can be found in **Table 1**. Prior to the fusion PCR step, DNA fragments located upstream and downstream to the epitope insertion sites were amplified by PCR using the Q5® High-Fidelity 2 × Master Mix (NEB) with relevant primers (**Table 1**) on a ProFlex PCR system (Life Technologies). After purification, fragments were ligated by performing a fusion PCR. The fused fragments were amplified by another PCR using the relevant external primer pairs (**Table 1**). Next, pBlueScript SK(+) harboring the p6- GLuc / p6 genomes or pTM-ORF1 vectors and epitope-tagged inserts were restricted by the suitable restriction enzymes (**Table 1**) and subsequently ligated using the T4 DNA ligase (NEB). Finally, *E. coli* (strain TOP10) were transformed with the ligated plasmid, selected clones were grown overnight at 37°C under agitation and the plasmid DNA was extracted and purified using the NucleoSpin plasmid Mini or Midi kit (Macherey-Nagel).

**Table 1.**
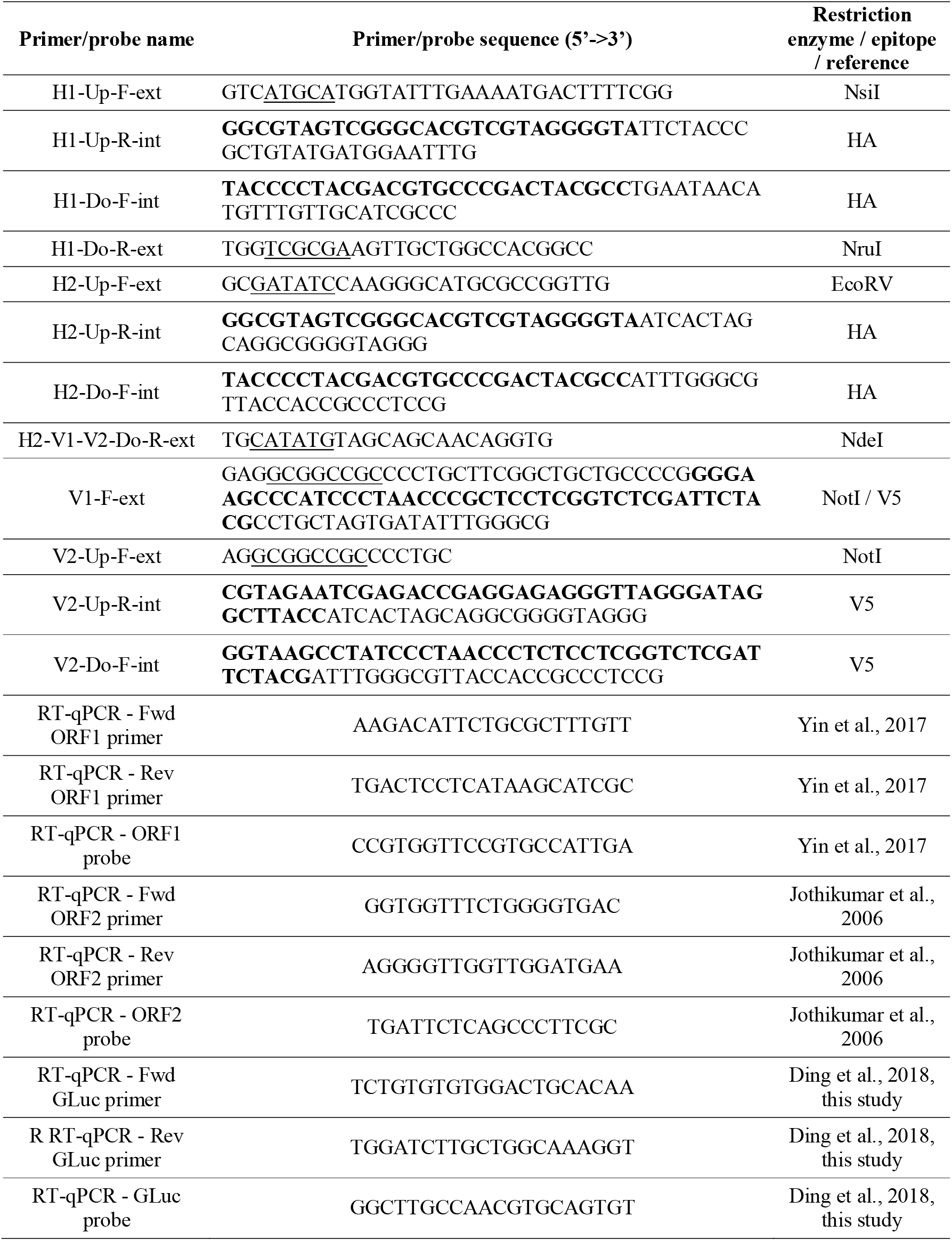
Sequences of primers and probes used to make the epitope tagged-ORF1 constructs and to quantify the genomic and subgenomic HEV RNAs by RT-qPCR. Each primer is named according to (i) the epitope insertion site (H1, H2, V1 or V2), (ii) upstream (Up) or downstream (Do) fragment, (iii) forward (F) or reverse (R) sense and (iv) external (ext) or internal (int) position. First, upstream and downstream fragments are amplified separately for each construct (*i.e.* for p6-H1-GLuc: upstream fragment amplified using H1-Up-F-ext / H1-Up-R-int and downstream fragment amplified using H1-Do-F-int / H1-Do-R-ext). Second, upstream and downstream fragments are fused thanks to the overlapping region. Third, the whole fragment containing the epitope is amplified using external primers (*i.e.* for p6-H1-GLuc: H1-Up-F-ext / H1-Do-R-ext). Note that the same primer H2-V1-V2-Do-R-ext was used for p6-H2-GLuc, p6-V1-GLuc and p6-V2-GLuc constructs. The p6-V1-GLuc construct was made using a single round of amplification and one primer pair: V1-F-ext / H2-V1-V2-Do-R-ext. Restriction sites are underlined in external primers. HA or V5 epitope sequence is highlighted in bold. Primers and probes used in RT-qPCR are listed. Fwd = forward, Rev = Reverse

### Capped mRNA synthesis

First, the plasmid DNA of p6-GLuc and p6 constructs were linearized using the MluI restriction enzyme (NEB). Next, the DNA was mixed with sodium acetate (3 M, pH 5.5) and chloroform /isoamyl alcohol (96v / 4v) and centrifuged at 14,000 rpm for 4 min. Following precipitation with absolute ethanol, the DNA was washed twice with 70% ethanol, dried and suspended in RNase free water. The capped mRNAs were synthetized by *in-vitro* transcription of the MluI-linearized DNA according to the mMESSAGE mMACHINE kit (Ambion) procedure and stored at -80°C before electroporation in PLC3 cells.

### Cell culture and transfection

PLC3 cells are a subclone of the PLC/PRF/5 (CRL-8024) hepatoma cells and were characterized as highly replicative and productive cell line for HEV (Montpellier et al., 2018). PLC3 cells were cultured in Dulbecco’s modified Eagle’s medium (DMEM, ThermoFisher Scientific) containing 10% inactivated fetal calf serum and 1% non-essential aa (ThermoFisher Scientific, DMEM complete).

Capped mRNA of either luciferase p6 replicons (p6-wt-GLuc, p6-GAD-GLuc mutant replicons and HA- or V5-ORF1-tagged p6-GLuc replicons, named as p6-H1-GLuc, p6-H2-Gluc, p6-V1-GLuc and p6-V2-GLuc) or p6 infectious strains (p6-wt and V5-tagged ORF1 p6 constructs named as p6-V1 and p6-V2) were electroporated in PLC3 cells as follows. After trypsinization, cells were suspended in DMEM complete medium and washed twice in Opti-MEM medium (ThermoFisher Scientific). Three million cells were electroporated with 10 µg of RNA and suspended in DMEM complete medium.

Huh-7.5 cells (RRID:CVCL_7927) are hepatoma cells derived from Huh-7 cells (Blight et al., 2002). They were grown and electroporated with the luciferase p6 replicons as described above.

Electroporated PLC3 or Huh-7.5 cells were treated with the HEV-replication inhibitor sofosbuvir (Selleck Chemicals) at 20 µM final concentration as previously published (DaoThi et al., 2016; Farhat et al., 2018).

Huh-7-derived H7-T7-IZ cells (Romero-Brey et al., 2012) stably expressing the T7 RNA polymerase were kindly provided by Dr R. Bartenschlager (University of Heidelberg, Germany). H7-T7-IZ cells were maintained in a DMEM completed medium supplemented with 50 µg/ml of Zeocin (InvivoGen) and used for transfection of the T7 promoter-driven pTM expression vectors. The pTM plasmids were transfected into H7-T7-IZ cells using TransIT®-LT1 Transfection Reagent (Mirus Bio LLC) following the manufacturer’s recommendations with a ratio ADN to transfection reagent of 1 to 3.

PLC3 cells were authenticated by STR profiling (Multiplexion). Huh-7.5 and H7-T7-IZ cells were authenticated by Multiplex Cell Authentication (Multiplexion).

### Luciferase assay

The electroporated cells were seeded in 24-well plates (80,000 cells/well) and incubated for 5 days at 37°C and 5% CO_2_. The supernatants were sampled at 1, 3, 4 and 5 days post-electroporation (dpe) and stored at -20°C until luminometer reading.

The supernatants were centrifuged at 14,000 rpm for 5 min and next, the samples were diluted 1:100 in 1 × passive lysis buffer (Promega) and 5 µl were transferred into a white Nunc 96-well plate. At 1 × s after injection of 20 µl of the substrate solution (Renilla Luciferase Assay System, Promega), relative light units (RLUs) were acquired on a Centro XS^3^ LB 960 luminometer (Berthold Technologies) during 10 s.

### Quantification of viral RNA

After electroporation of PLC3 cells with the p6 luciferase replicons or the infectious p6 strains, total RNA was extracted from cellular supernatants (QIAamp Viral RNA Mini kit, Qiagen) or cells (NucleoSpin RNA plus kit, Macherey-Nagel) at different time post-electroporation (4 hpe – 7 dpe). The RNA was next converted to cDNA by using a polydT primer and following the AffinityScript Multiple Temperature cDNA Synthesis kit instructions (Agilent Technologies). In order to quantify the HEV genome, a standard curve was prepared by diluting the *in vitro*-transcribed HEV p6 plasmid in total RNA extracted from mock electroporated PLC3 cells. For the quantification of intra- and extra-cellular HEV RNA, primers and probes were designed against genomic and subgenomic RNAs according to previously published literature (**Table 1**) (Jothikumar et al., 2006; Yin et al., 2017; Ding et al., 2018). The viral RNA copy numbers were quantified by qPCR (TaqMan Gene Expression Assay, MGB-FAM-dye, ThermoFisher Scientific) using the QuantStudio3 Thermocycler (Applied Biosystems).

### Cell lysis, subcellular fractionation and immunoblotting

The cells were lysed in B1 buffer (100 mM NaCl, 2 mM EDTA, 1% Triton-X100, 50 mM Tris-HCl, 0.1% SDS) including 1 mM PMSF and 1×protease inhibitors (cOmplete^TM^ protease inhibitor cocktail, Roche) and stored at -20°C in 1 Laemmli buffer until usage. Subcellular fractionation was performed using the subcellular protein fractionation kit for cultured cells following the supplier instructions (ThermoFisher Scientific).

The non-infectious samples were heated at 70°C for 10 min while the infectious samples were inactivated at 80°C for 20 min. Samples were then loaded on an 8% SDS-PAGE gel and then transferred onto a nitrocellulose membrane (Hybond-ECL, Amersham). The membrane was incubated in blocking buffer (1 × PBS with 5% milk and 0.2% Tween-20) for 1 h at RT under constant shaking. The primary antibody (**Table 2**) was incubated overnight at 4°C under constant shaking in 1 × PBS containing 0.2% Tween-20 and 2% BSA. The membrane was washed 3 times with a solution of 1 × PBS and 0.2% Tween-20, which was followed by an incubation with a suited peroxidase-conjugated secondary antibody in blocking buffer for 45 min at RT. The membrane was washed again 3 times. Finally, the proteins were detected by chemiluminescence using the Pierce ECL Western Blotting Substrate (Life Technologies) and a developer.

**Table 2.**
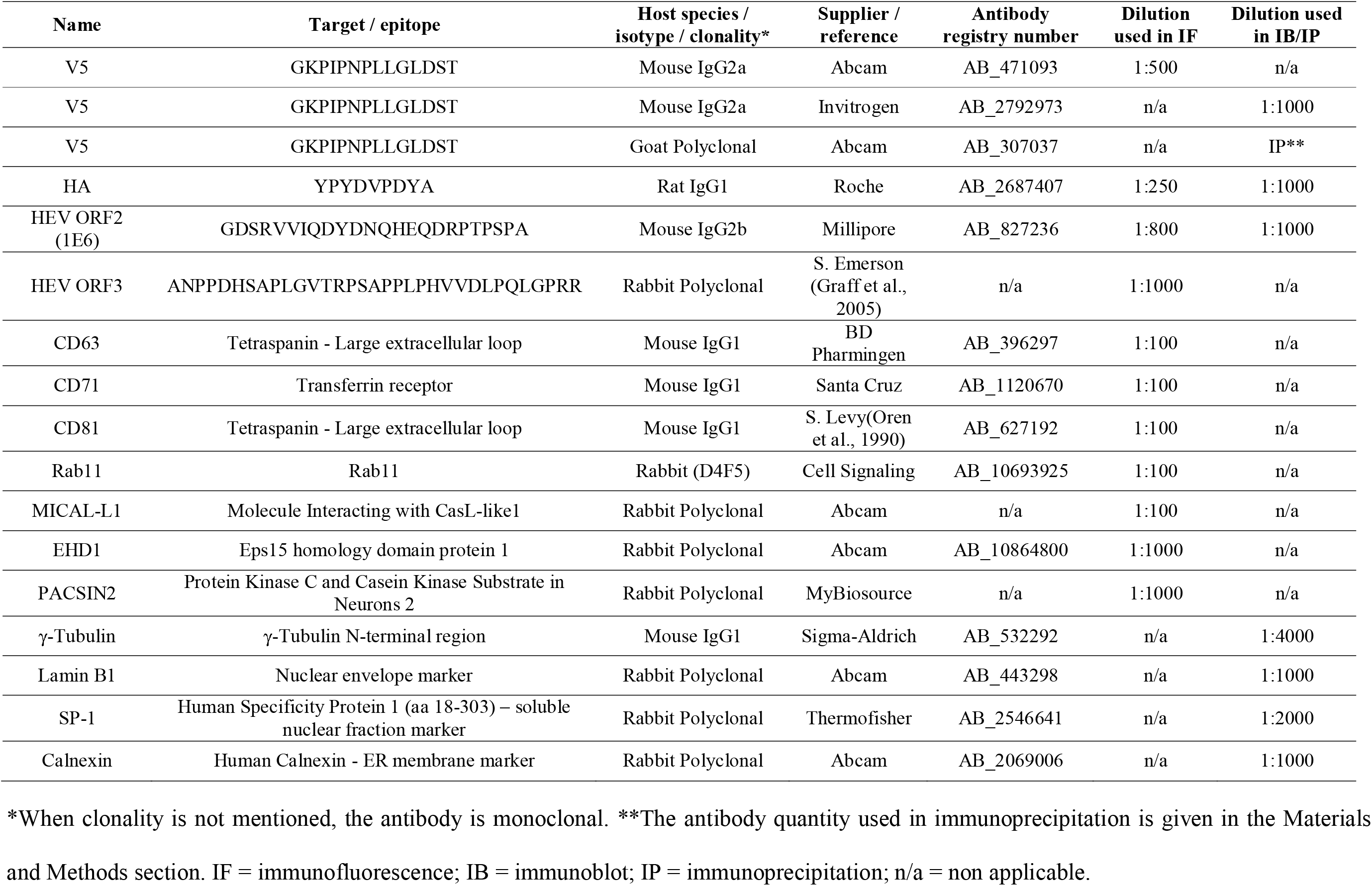
List of primary antibodies used in this study.

### Infectious Titers

After electroporation of PLC3 cells with p6-wt, p6-V1 or p6-V2 RNA or PBS, 1 × 10^6^ cells were seeded into T-75 flasks in DMEM complete medium. Eight hours after seeding, the medium was changed to HEV medium: DMEM/M199 (1v:1v), 1 mg/ml of lipid-rich albumin (AlbuMAX^TM^ I Lipid-rich BSA, ThermoFisher Scientific), 1% non-essential aa and 1% pyruvate sodium (ThermoFisher Scientific). Then, the HEV producing cells were incubated at 32°C and 5% CO_2_ for 10 days.

Next, Huh-7.5 cells seeded into coated 96-well plates (8,000 cells/well) were infected with undiluted and serially diluted supernatants from HEV-producing cells. The inoculum was removed after 8 h and cells were overlaid with fresh medium. Three days post-infection at 37°C and 5% CO_2_, cells were fixed and processed for indirect immunofluorescence with anti-ORF2 antibody 1E6 (Millipore). For confocal microscopy analysis, Huh-7.5 cells were seeded on glass coverslips in 24-well plates and infected with 500 µl of undiluted infectious cell supernatant.

### Indirect immunofluorescence

Cells were first fixed with 3% paraformaldehyde for 20 min, washed three times with 1 × PBS, then permeabilized for 5 min with cold methanol and subsequently with 0.5% Triton X-100 for 30 min. Cells were incubated with 10% goat serum diluted in 1 × PBS for 30 min at RT and then stained with primary antibodies (**Table 2**) for 30 min at RT followed by three washes with 1 × PBS and then incubated with a suited secondary antibodies for 20 min at RT. DAPI (4’,6-diamidino-2-phenylindole, 1:500) was used to stain the nuclei. Finally, coverslips were mounted with Mowiol®4-88 (Calbiochem) on glass slides. Cells were analyzed using an LSM880 confocal laser-scanning microscope (Zeiss) using a ×63/1.4 numerical aperture oil immersion lens. The images were next processed using the ImageJ and Fiji softwares. Colocalization studies were performed by calculating the Pearson’s Correlation Coefficient (PCC) using the JACoP plugin of ImageJ and Fiji softwares. The PCC examines the relationship between the intensities of pixels from two channels in the same image. Twenty-to-thirty whole cells or regions of interest (ROI) were analyzed to calculate the PCC mean. A PCC of 1 indicates perfect correlation, 0 no correlation, and -1 a perfect anti-correlation.

### Immunoprecipitations

Immunoprecipitations (IP) were performed with cell lysates of either electroporated PLC3 cells (3 dpe) or transfected H7-T7-IZ cells (1 dpt). Protein G sepharose beads (GE Healthcare, 40 µl per IP) were equilibrated in B1 lysis buffer by washing and centrifuging them at 6,000 rpm at 4°C. The beads were then incubated with the anti-V5 goat polyclonal antibody (Abcam, 1 μg of anti-V5 per mg of total proteins) in B1 buffer overnight at 4°C on a spinning wheel. The cell lysates were pre-cleared with 40 µl of beads without specific antibodies for 30 min on a spinning wheel. Next, the pre-cleared cell lysates were added to the antibody-bound beads for 2 h at RT. After this, the beads were washed 6 times with 1 × PBS containing 0.5% NP-40. The samples were heated for 10 min at 70°C or inactivated at 80°C for 20 min and loaded on an 8% SDS-PAGE gel.

### Mass spectrometry analysis

Proteins were resolved by SDS-PAGE. Colloïdal blue stained bands corresponding to ORF1 proteins in WB were cut for in-gel digestion with trypsin. NanoLC-MS/MS analyses of the protein digests were performed on an UltiMate-3000 RSLCnano System coupled to a Q-Exactive instrument (Thermo Fisher Scientific), as previously described (Montpellier et al., 2018). Collected raw data were processed and converted into *.mgf peak list format with Proteome Discoverer 1.4 (Thermo Fisher Scientific). MS/MS data were interpreted using search engine Mascot (version 2.4.0, Matrix Science) with a tolerance on mass measurement of 0.2 Da for precursor and 0.2 Da for fragment ions, against a composite target decoy database (40584 total entries) built with Swissprot *Homo sapiens* database (TaxID=9606, 20 May 2016, 20209 entries) fused with the sequences of ORF1 (p6_AFD33683) and a list of classical contaminants (119 entries). Carbamidomethylation of cysteine residues, oxidation of methionine residues, protein N-terminal acetylation and propionamidation of cysteine residues were searched as variable modifications. Up to one trypsin missed cleavage was allowed. Semi-specific cleavage was also authorized.

### *In situ* labeling of viral RNA

The RNAscope® is a branched DNA *in situ* hybridization technology that specifically and sensitively detects RNA in fixed cells or tissues. First, 3 pools of dual Z-shaped probes of 18-25 nucleotides were designed against the HEV genomic and subgenomic RNAs (Advanced Cell Diagnostics Bio-Techne, ACD Bio, **Figure 8A**). Two probes need to bind next to each other to produce a specific signal. Each target probe set contains a pool of 20 oligo ZZ pairs. PLC3 cells electroporated with p6-wt or p6-V1 infectious strains were fixed in 3% PFA for 20 min. Coverslips holding the fixed cells were attached to glass slides with a drop of nail polish and hydrophobic barriers were drawn around them with the ImmEdge Hydrophobic Barrier Pen (ACD Bio). Next, fixed cells were pre-treated according to the supplier instructions (RNAscope® H_2_O_2_ and Protease Reagents). First, the cells were treated with H_2_O_2_ for 10 min at RT and then washed twice with 1 × PBS. Next, the protease III was diluted 1:15 and incubated for 15 min at RT; slides were washed twice. Then, the RNAscope assay was carried out following the user manual precisely (RNAscope Detection Kit Multiplex Fluorescent Reagent Kit v2 (Wang et al., 2012; Liu et al., 2019). The positive and negative strand of genomic viral RNAs were targeted by probe A (ref. 1030631-C2, ACD Bio) and probe B (ref. 579841-C3, ACD Bio), respectively (**Figure 8A**). Probe C (ref. 586651-C1, ACD Bio) targeted the positive strand of subgenomic RNA (**Figure 8A**). The RNAs were labeled using the Opal 520, Opal 570 and Opal 650 as fluorophores (Akoya Biosciences). Subsequently, immunofluorescent labeling using either the anti-V5, 1E6 or anti-ORF3 antibodies could be performed (**Table 2**). Finally, coverslips were mounted and cells were analyzed by confocal microscopy as mentioned above.

### Statistical analysis

Mann-Whitney statistical tests were performed with R version 3.6.1. Any test with a pvalue < 0.05 is declared as significant.

## Results

### Insertion of epitope tags in the ORF1 polyprotein of HEV

Since there is no commercial antibody that recognizes the ORF1 protein in HEV-replicating cells, we first sought to insert epitope tags in the ORF1 sequence in a way to preserve the HEV replication and to allow the characterization of the HEV replicase. We aligned 44 aa sequences from different HEV genotypes to identify variable regions which are generally more prone to accept epitope insertions without impacting viral replication. Indeed, the HVR stood out as the region with the highest divergence of the entire HEV genome, thus confirming previous reports (Muñoz-Chimeno et al., 2020). Focusing on the HVR, we next aligned several epitope aa sequences with that of HEV in an aim to identify similar aa and to ease epitope insertion with the least disruption. Such a location highly similar to the V5 epitope aa sequence was found in the HVR (**Figure 1C**). While 4 aa residues remained unchanged (G729, P731, P733, P735), 3 were mutated (L730K, H732I, T734N) and 7 were inserted to generate the full V5 epitope (V1, **Figure 1C**). In a second approach, we took advantage of the report by Nguyen et al. to insert a V5 and a HA epitope tags at position 2143 (V2, H2, **Figure 1C**) (Nguyen et al., 2012). These authors isolated a HEV gt3 strain from a chronically infected patient in which the human S19 ribosomal coding sequence was inserted at that corresponding position and led to a HEV replication advantage in cell culture (Nguyen et al., 2012). Finally, a HA tag was also introduced at the C-terminus of the RdRp, since avoiding an insertion within the ORF1 protein was supposed to maintain viral replication (H1, **Figure 1C**).

### Replication of HEV p6 replicons expressing epitope-tagged ORF1

To evaluate whether epitope insertions modify the replication efficacies, we made use of the HEV *Gaussia* luciferase replicon (GLuc) in which the *GLuc* reporter gene is transcribed by the ORF1 viral replicase (**Figure 1B**, (Shukla et al., 2012; Emerson et al., 2013). The luciferase, secreted in cell supernatants, is used as a readout of ORF1 replication efficacy. To measure the impact of V5 and HA epitope insertion on HEV replication, we electroporated the *GLuc* replicons expressing wt ORF1 (p6-wt-GLuc), polymerase-inactivated GAD mutant (p6-GAD-GLuc), HA-tagged ORF1 (p6-H1-GLuc, p6-H2-GLuc) or V5-tagged ORF1 (p6-V1-GLuc, p6-V2-GLuc) in PLC3 and Huh-7.5 cells, which have been described as efficient HEV cell culture systems (Farhat et al., 2018; Montpellier et al., 2018). The replication efficacies were analyzed over a course of 5 days post-electroporation (dpe) and replication folds (normalized to 1 dpe) were compared (**Figures 2A,B**). The p6-H1-GLuc and p6-H2-GLuc replicons respectively showed a 95% and 80% reduction of replication efficacies at 5 dpe, as compared to the p6-wt-GLuc replicon in PLC3 cells. In Huh-7.5 cells, the replications efficacies of both constructs were also drastically reduced (97% and 76% reduction for p6-H1-GLuc and p6-H2-GLuc, respectively). Due to poor replication efficacies, these constructs were not further characterized. In contrast, the replication kinetics of the p6-V1-GLuc and p6-V2-GLuc replicons were similar to that of p6-wt-GLuc, in both PLC3 and Huh-7.5 cell lines (**Figures 2A,B**). Thus, the V5 epitope insertions at the selected positions in the ORF1 HVR do not alter HEV replication.

**Figure 2.**
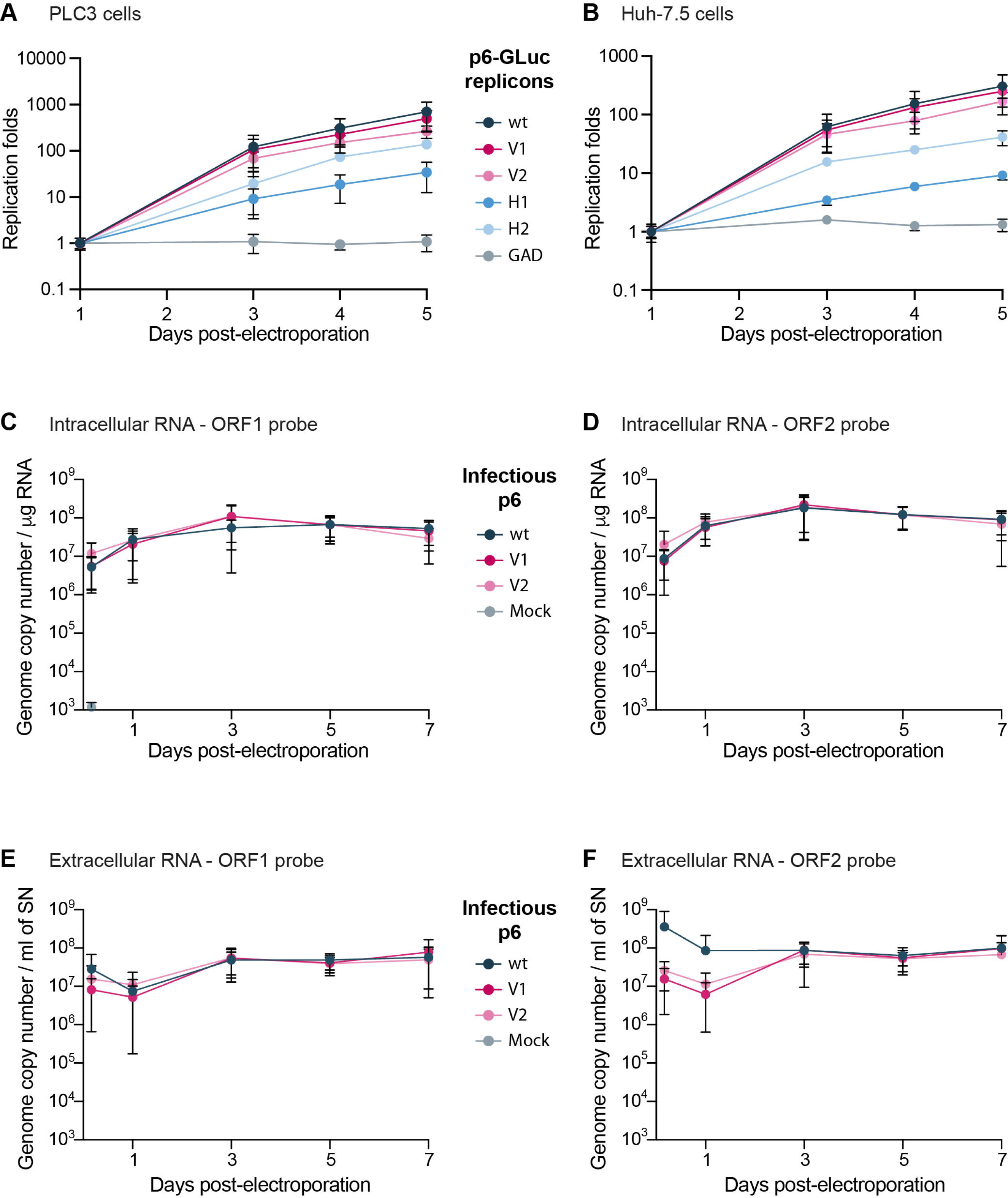
Replication efficacies of HEV p6 replicons and infectious strain expressing epitope-tagged ORF1. **Figures 2A, 2B.** Replication efficacies of HEV replicons expressing tagged ORF1 in PLC3 (**Figure 2A**) and Huh-7.5 cells (**Figure 2B**) as measured by luciferase activity. The relative light units (RLU) were measured everyday for 5 days post-electroporation (dpe) by quantification of the *Gaussia* luciferase in the cell supernatants using a luminometer. Replication fold increases of the tagged p6-GLuc replicons (V1, V2, H1, H2) and non-tagged p6-GLuc replicons (wt, GAD) were normalized to 1 dpe. The p6-GAD-GLuc is a non-replicative construct that possesses a mutation from GDD to GAD which inactivates the ORF1 polymerase. Experiments were conducted three times with 3 technical replicates. **Figures 2C-F.** HEV RNA quantification in p6 electroporated-PLC3 cells expressing wild type or epitope-tagged ORF1. Intracellular (**Figures 2C,D**) and extracellular (**Figures 2E,F**) viral RNAs were quantified at 4 h post-electroporation (hpe), 1, 3, 5 and 7 dpe by RT-qPCR targeting either ORF1 (**Figures 2C,E**) or ORF2 (**Figures 2D,F**). Mock-electroporated cells were used as negative controls and fluorescence signal was at or below the detection limit. Experiments were conducted three times with 2 technical replicates.

Of note, in the presence of sofosbuvir (20 µM), a nucleotide analogue that efficiently inhibits HCV and HEV polymerases (DaoThi et al., 2016; Farhat et al., 2018), the replication efficacies of all replicons were inhibited at 5 dpe by 76 to 87% in PLC3 cells and by 73% to 92% in Huh-7.5 cells (**Supplementary Figures 1A,B**). These results indicate that the insertion of epitope tags in the ORF1 does not affect the sofosbuvir efficacy to inhibit HEV replication.

### Replication of HEV infectious genome expressing epitope-tagged ORF1

The V5 epitope was next introduced into the p6 infectious full-length genome at 2 positions, leading to the p6-V1 and p6-V2 constructs (**Figure 1C**). Intra- and extracellular HEV RNA levels were measured from 4 hours post-electroporation (4 hpe) to 7 dpe, by RT-qPCR using probes against the genomic RNA (ORF1 probe, **Figure 1A**, **Table 1**) or the genomic and subgenomic RNAs (ORF2 probe, **Figure 1A**, **Table 1**). The intracellular RNA levels increased rapidly for p6-V1 and p6-V2 constructs within 1 dpe to reach approximately 1x10^8^ copies / μg RNA (ORF1 probe, **Figure 2C**) and 2x10^8^ copies / μg RNA (ORF2 probe, **Figure 2D**) at 3 dpe. Then, the RNA levels decreased slightly but remained above 3x10^7^ (ORF1 probe, **Figure 2C**) and 7x10^7^ (ORF2 probe, **Figure 2D**) copies / μg RNA at 7 dpe. The intracellular RNA copy numbers of the p6-V1 and p6-V2 constructs followed similar kinetics to that of p6-wt from 4 hpe to 7 dpe. In the cell supernatants, all RNA copy numbers decreased during the first day, then increased to reach a level of 5.5x10^7^ (ORF1 probe, **Figure 2E**) and 9x10^7^ (ORF2 probe, **Figure 2F**) copies / μg RNA at 3 dpe and remained constant towards the end of the experiment. The RNA levels measured for the p6-V1 and p6-V2 constructs in the cell supernatants were comparable to the p6-wt RNA level evolution. Moreover, to check whether ORF2 expression from subgenomic RNA was not altered, PLC3 cells electroporated with the p6-V1 or p6-V2 constructs were processed for ORF2 immunofluorescence (**Supplementary Figure 1C**). V5-tagged-HEV-producing cells displayed an ORF2 fluorescent labeling similar to that of p6-wt-electroporated cells. Additionally, the expression of ORF2 was inhibited by sofosbuvir as a much lower fluorescent signal could be visualized in the treated electroporated cells compared to the non-treated cells (**Supplementary Figure 1C**). Taken together, these results indicate that the insertions of V5 epitopes at the selected positions in ORF1 HVR do not affect replication efficacy of the p6 genome in PLC3 cells.

### Expression and processing of the V5-tagged HEV replicase

We next sought to analyze the expression and processing of the ORF1 polyprotein in HEV replicative systems. Since the ORF1 protein has been largely studied in heterologous expression systems, we compared the expression and maturation of V5-tagged ORF1 in replicative and heterologous systems. For this purpose, the V5-tagged ORF1 were cloned downstream of the T7 promoter into the pTM plasmid and expressed in H7-T7-IZ cells, as previously described (Lenggenhager et al., 2017).

First, the expression of the V5-tagged ORF1 was analyzed from 8 hours (8 hpt) to 3 days post-transfection (3 dpt) in the H7-T7-IZ heterologous expression system. As early as 8 hpt, the ORF1 protein was detectable by anti-V5 immunoblot as a major band migrating above 180 kDa, a size that corresponds to the expected molecular weight of the full-length ORF1 (**Figure 3A**). This major band was present at every time points and was accompanied by several lower molecular weight bands of lesser intensity, ranging from 95 to 170 kDa, which may correspond to ORF1 cleavage products (**Figure 3A**, stars). Second, the expression kinetics of the V5-tagged ORF1 protein was monitored by immunoblot, from 4 hpe to 3 dpe, in PLC3 cells electroporated with the p6-wt-GLuc, p6-V1-GLuc and p6-V2-GLuc replicons (**Figure 3B**). As early as 4 hpe, a band of high signal intensity was detected above 180 kDa, which corresponds to the full-length ORF1 protein. Additionally, several bands of lower molecular weights (95 to 170 kDa) were also detected and may correspond to potential cleavage products of the ORF1 polyprotein. At 3 dpe, the overall smaller-size band pattern ranging from 95 kDa to 170kDa was comparable to the band profile observed at early time points. At last, the expression profiles of the V5-tagged ORF1 in PLC3 electroporated with the p6 infectious strains (p6-wt, p6-V1, p6-V2, **Figure 3C**) were compared to the profiles of ORF1 expressed in the replicon and heterologous systems (**Figures 3A,B**, respectively). Similarly, the most intense band was migrating above 180 kDa at every time points and smaller bands were identified between 95-180 kDa. The V5-tagged ORF1 expression signal was more intense at early time points (4 hpe-3 dpe) when compared to later time points (15 and 25 dpe, **Figure 3C**). The V5-tagged ORF1 expression level decreased but was still detectable at 15 and 25 dpe, as it was for the ORF2 capsid protein, thus showing that the HEV expressing a V5-tagged replicase is able to fulfill its infectious cycle in the long-term. Thus, the successful expression of ORF1 polyprotein allowed the detection of an abundant full-length protein but also of less abundant smaller size products in the 3 systems analyzed. To exclude that the V5-tagged ORF1 minor bands were due to protein degradation, proteasome inhibition experiments were conducted (**Supplementary Figure 2**). Upon treatment with lactacystin (30 μM for 8 hours), the V5-tagged ORF1 minor bands were still detectable in both H7-T7-IZ and PLC3 cells and their pattern remained unchanged. As a control, we probed for HSP70 protein that accumulated in the lactacystin-treated conditions, indicating that the proteasome inhibition was successful (Liao et al., 2006; Young and Heikkila, 2010). Thus, these minor ORF1 bands are likely not the result of proteasome degradation.

**Figure 3.**
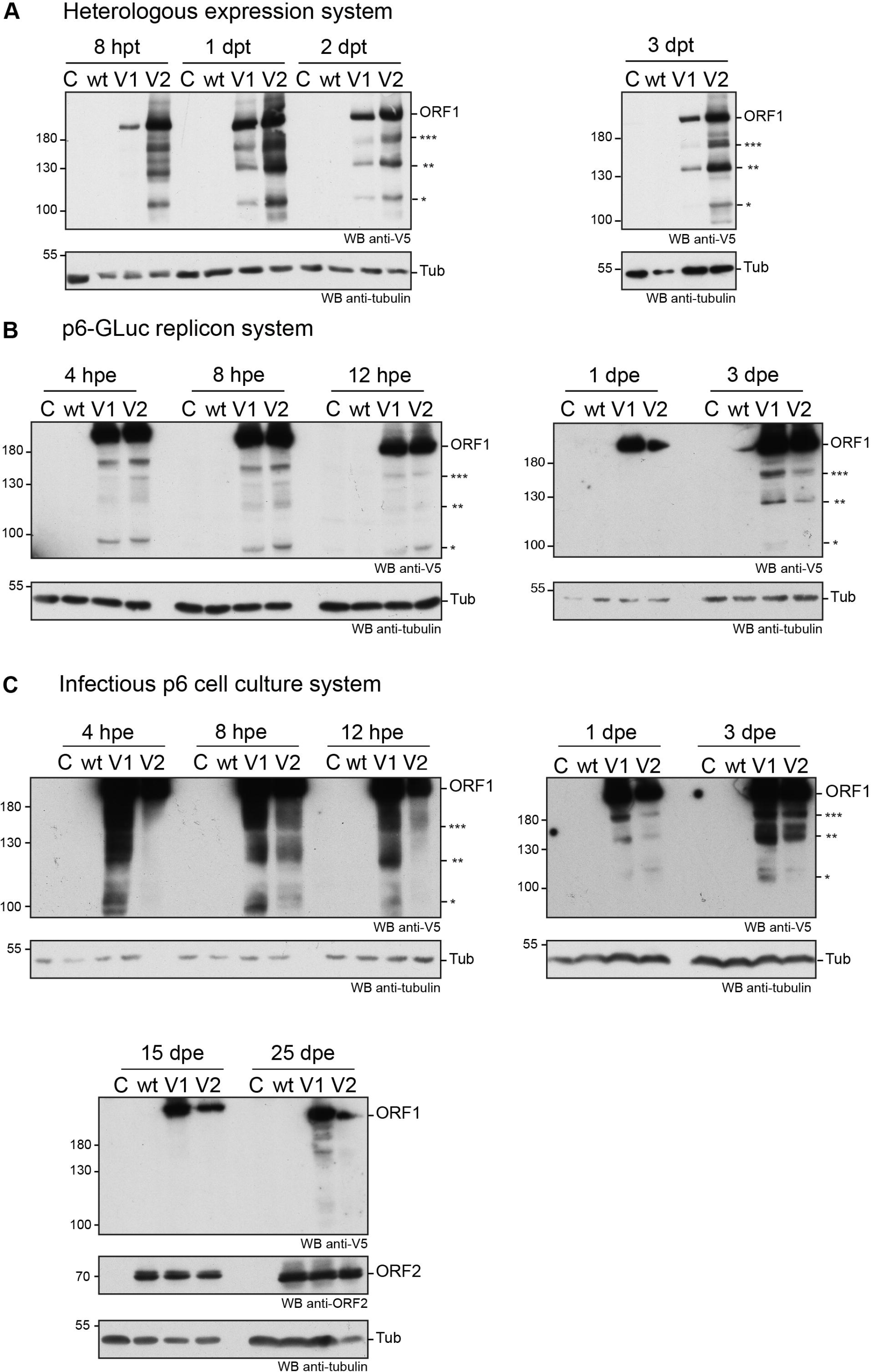
Expression of the V5-tagged ORF1 protein over time in different cellular systems. **Figure 3A.** Heterologous expression of the HEV p6 ORF1 protein. H7-T7-IZ cells were transfected with a pTM plasmid expressing the wild type untagged ORF1 protein (wt) or the V5-tagged ORF1 (V1 or V2). The H7-T7-IZ cells constitutively express the T7 polymerase and the ORF1 gene lies under the control of a T7 promoter. Total cell lysates were collected in presence of protease inhibitors at 8 hours post-transfection (hpt) and 1, 2 and 3 days post-transfection (dpt). Mock-transfected cells served as negative control (C). The band migrating higher than 180 kDa corresponds to the full-length ORF1 protein (ORF1). The immunoblot was probed either with an anti-V5 antibody or an anti-γ-tubulin antibody to control for even loading. Molecular weight markers are indicated in kilodaltons. Stars indicate lower molecular weight ORF1 products. **Figure 3B.** Expression of the ORF1 protein in the p6-GLuc replicon system. PLC3 cells were electroporated with a replicon expressing the wild type untagged ORF1 protein (wt) or the V5-tagged ORF1 (V1 or V2). Total cell lysates were collected in presence of protease inhibitors at 4, 8, 12 hpe and 1 and 3 dpe. Mock-electroporated cells served as negative control (C). The band migrating higher than 180 kDa corresponds to the full-length ORF1 protein (ORF1). The immunoblot was probed either with an anti-V5 antibody or an anti-γ-tubulin antibody to control for even loading. Molecular weight markers are indicated in kilodaltons. Stars indicate lower molecular weight ORF1 products. **Figure 3C.** Expression of the ORF1 protein in the infectious p6 cell culture system. PLC3 cells were electroporated with *in vitro* transcribed genomic RNA of the p6 infectious strain expressing the wild type untagged ORF1 protein (wt) or the V5-tagged ORF1 (V1 or V2). Total cell lysates were collected in presence of protease inhibitors at 4, 8, 12 hpe and 1, 3, 15, 25 dpe. Mock-electroporated cells served as negative control (C). The band migrating higher than 180 kDa corresponds to the full-length ORF1 protein (ORF1). The immunoblot was probed either with an anti-V5 antibody or an anti-ORF2 (1E6) antibody or an anti-γ-tubulin antibody to control for even loading. Molecular weight markers are indicated in kilodaltons. Stars indicate lower molecular weight ORF1 products.

In an aim to identify these potential ORF1 cleavage products, V5-immunoprecipitations were set up from cell lysates produced from all 3 systems (**Figures 4A,B**). The pattern of the smaller-size ORF1 proteins differed from one expression system to another with some proteins migrating at the same size (**Figures 4A,B**, stars). Minor ORF1 bands appeared more numerous in the p6 infectious cell culture system than in other systems. As better ORF1 expression levels were achieved in both the heterologous and p6 cell culture systems, V5-immunoprecipitations were scaled up from these 2 systems and trypsin-digests of some ORF1 selected products were analyzed by nano-scale liquid chromatography coupled to tandem mass spectrometry (**Figures 4C,D**, **Table 3**). The full-length V5-tagged ORF1 proteins, expressed in the heterologous and infectious p6 cell culture systems, were identified with 35% and 38% of peptide coverage, respectively. In spite of several attempts to reach sufficiently pure tagged-ORF1 protein in high-enough quantity, the peptide coverage of smaller size ORF1 proteins was not sufficient to identify any potential cleavage sites. Overall, the N-terminus of the V5-tagged ORF1 protein bands was better covered than the C-terminus, suggesting a limited processing at the polyprotein N-terminus especially in the heterologous system. Some higher molecular weight products (bands 8-10, **Table 3**), visible on the Coomassie-stained gel, but not detected by the V5 antibody, could correspond to ORF1 oligomers (**Figures 4C,D**).

**Figure 4.**
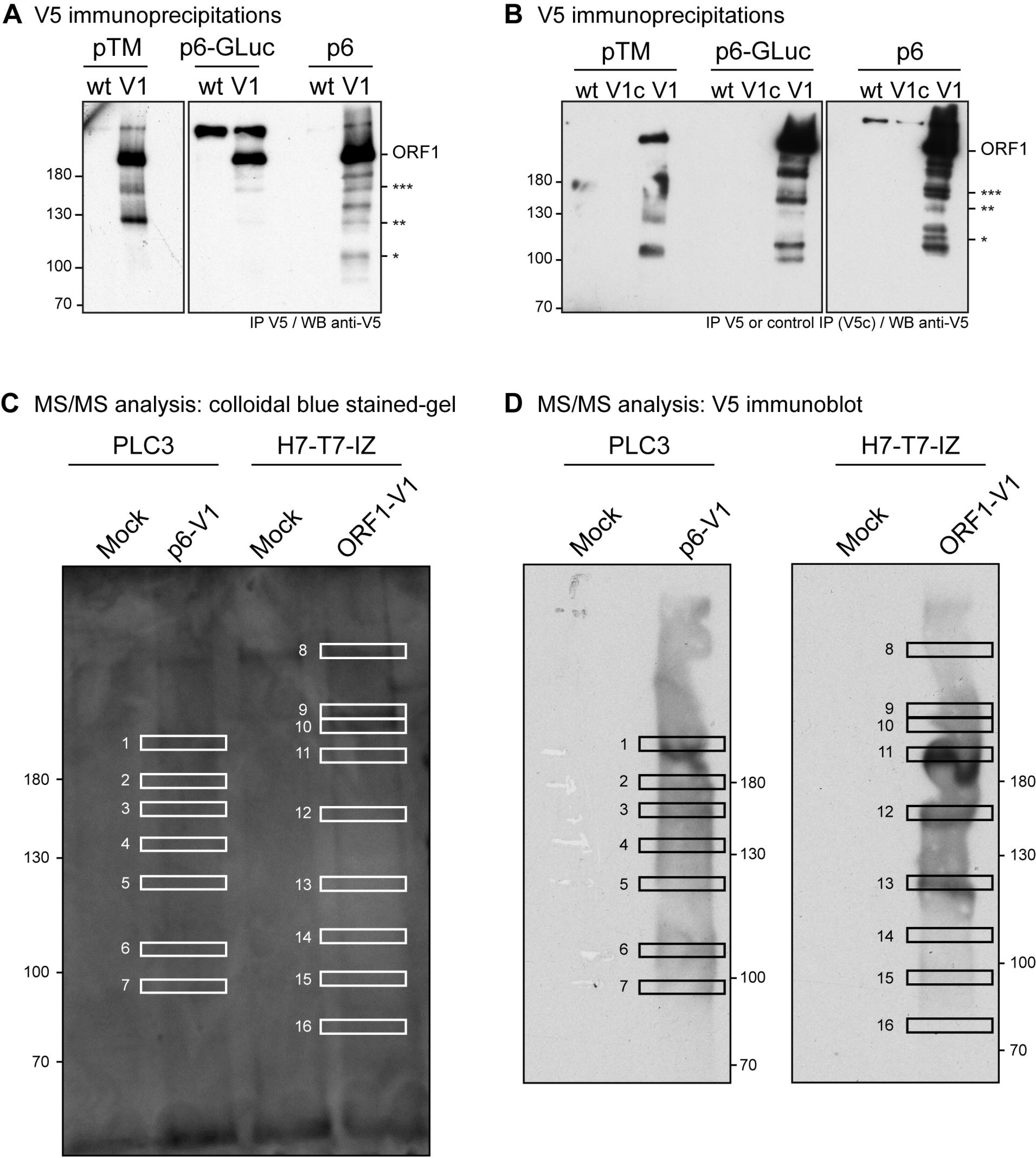
Immunoprecipitation of the V5-tagged ORF1 protein expressed in different cellular systems and mass spectrometry analysis. **Figure 4A.** HEV p6 ORF1 protein was heterologously expressed in H7-T7-IZ cells following transfection with the pTM plasmid carrying the untagged (wt) or V5-tagged (V1) construct (left panel). One dpt, total cell lysates were immunoprecipitated with the polyclonal goat anti-V5 antibody. HEV p6 protein was expressed in the replicon (p6-GLuc) and p6 cell culture (p6) systems (right panel). Three dpe, PLC3 cells lysates expressing either the untagged (wt) or the V5-tagged (V1) ORF1 proteins were immunoprecipitated using the polyclonal goat anti-V5 antibody. Immunoblots were revealed with a mouse anti-V5 monoclonal antibody. Molecular weight markers are indicated in kilodaltons. Stars indicate lower molecular weight ORF1 products. **Figure 4B.** In the same conditions as above, the control immunoprecipitations were conducted on the lysates expressing the V5-tagged ORF1 in all 3 systems using protein G sepharose beads without antibody (lanes V1c). The other lanes are labeled as in **Figure 4A.** Molecular weight markers are indicated in kilodaltons. Stars indicate lower molecular weight ORF1 products. **Figures 4C,D.** Immunoprecipitation of V5-tagged HEV replicase for mass spectrometry analysis. PLC3 cells were electroporated with the p6-V1 construct expressing a V5-tagged ORF1 (p6-V1). H7-T7-IZ cells were transfected with the pTM plasmid expressing ORF1-V1. Mock-transfected or -electroporated cells were used as negative controls (Mock). Cellular lysates were immunoprecipitated with the V5 antibody. Next, the eluate was partitioned in 2 fractions and resolved by SDS-PAGE electrophoresis. Most of the eluate was stained with colloidal blue (**Figure 4C**) while the remaining was probed with the anti-V5 antibody (**Figure 4D**). Following precise alignment of the WB and colloidal blue stained gel, the indicated bands (white rectangles) were cut from the gel and analyzed by nano-scale liquid chromatography coupled to tandem mass spectrometry after in-gel trypsin digestion (**Table 3**).

**Table 3.** Mass spectrometry analysis.

| **Band number / construct** | **Coverage (%)** | **Number of identified spectra** | **N-term. identified peptides (NSI)** | **C-term. identified peptides (NSI)** |
| --- | --- | --- | --- | --- |
| 1 / p6-V1 | 38 | 129 | 1-8 (2) | 1758-1777(1) |
| 2 / p6-V1 | 32 | 87 | 41-53(1) | 1720-1734(1) |
| 3 / p6-V1 | 31 | 68 | 86-94(1) | 1720-1734(1) |
| 4 / p6-V1 | 32 | 83 | 41-53(2) | 1695-1706(1) |
| 5 / p6-V1 | 30 | 68 | 41-53(1) | 1720-1734(1) |
| 6 / p6-V1 | 20 | 48 | 41-53(1) | 1651-1662(1) |
| 7 / p6-V1 | 18 | 41 | 86-94(2) | 1403-1414(1) |
| 8 / ORF1-V1 | 39 | 125 | 41-53(2) | 1720-1734(1) |
| 9 / ORF1-V1 | 55 | 321 | 1-8(3) | 1758-1777(3) |
| 10 / ORF1-V1 | 71 | 672 | 1-8(4) | 1758-1777(7) |
| 11 / ORF1-V1 | 35 | 195 | 1-8(10) | 1405-1419(2) |
| 12 / ORF1-V1 | 40 | 185 | 1-8(2) | 1651-1662(1) |
| 13 / ORF1-V1 | 31 | 102 | 1-8(2) | 1695-1706(1) |
| 14 / ORF1-V1 | 19 | 75 | 1-8(2) | 1651-1662(1) |
| 15 / ORF1-V1 | 17 | 63 | 1-8(2) | 1390-1398(1) |
| 16 / ORF1-V1 | 17 | 66 | 1-8(2) | 1390-1398(1) |

2 The full-length ORF1-V1 of the p6 Kernow C-1 strain is 1779 aa in length. Bands 8-10 were 3 visible on the colloidal blue stained gel but were not detected by the V5 antibody. NSI = 4 number of spectrums identifying peptide. Peptides identified only once with a score lower 5 than 25 were not taken into account.

We next performed subcellular fractionation of V5-tagged ORF1 expressing cells (**Figures 5A,B**). The V5-tagged ORF1 full-length protein (> 180 kDa) was detected by immunoblot in the soluble and membranous cytoplasmic fractions (Cs, Cm) as well as in the nuclear soluble fraction (Ns) and the nuclear envelope fraction (Ne) of p6-GLuc and p6 expression systems. The nuclear chromatin-bound fraction (Nc) only displayed a weak V5 signal in both expression systems. Also, smaller bands, migrating between 100 to 180 kDa (already detected in **Figure 3**), that could correspond to potential cleavage products of the ORF1 polyprotein, were detected in all fractions except in the Nc fraction. Notably, a subtle change in the band pattern was visible in the V5-tagged ORF1 expressed in PLC3 electroporated with the p6-V1: a signal was detected at 160 kDa in the Ns and Ne fractions (**Figure 5B**, 3 stars). This band was not detected either the Cs or the Cm fractions. Together, these results suggest that the ORF1 protein is likely partitioned in different cell compartments.

**Figure 5.**
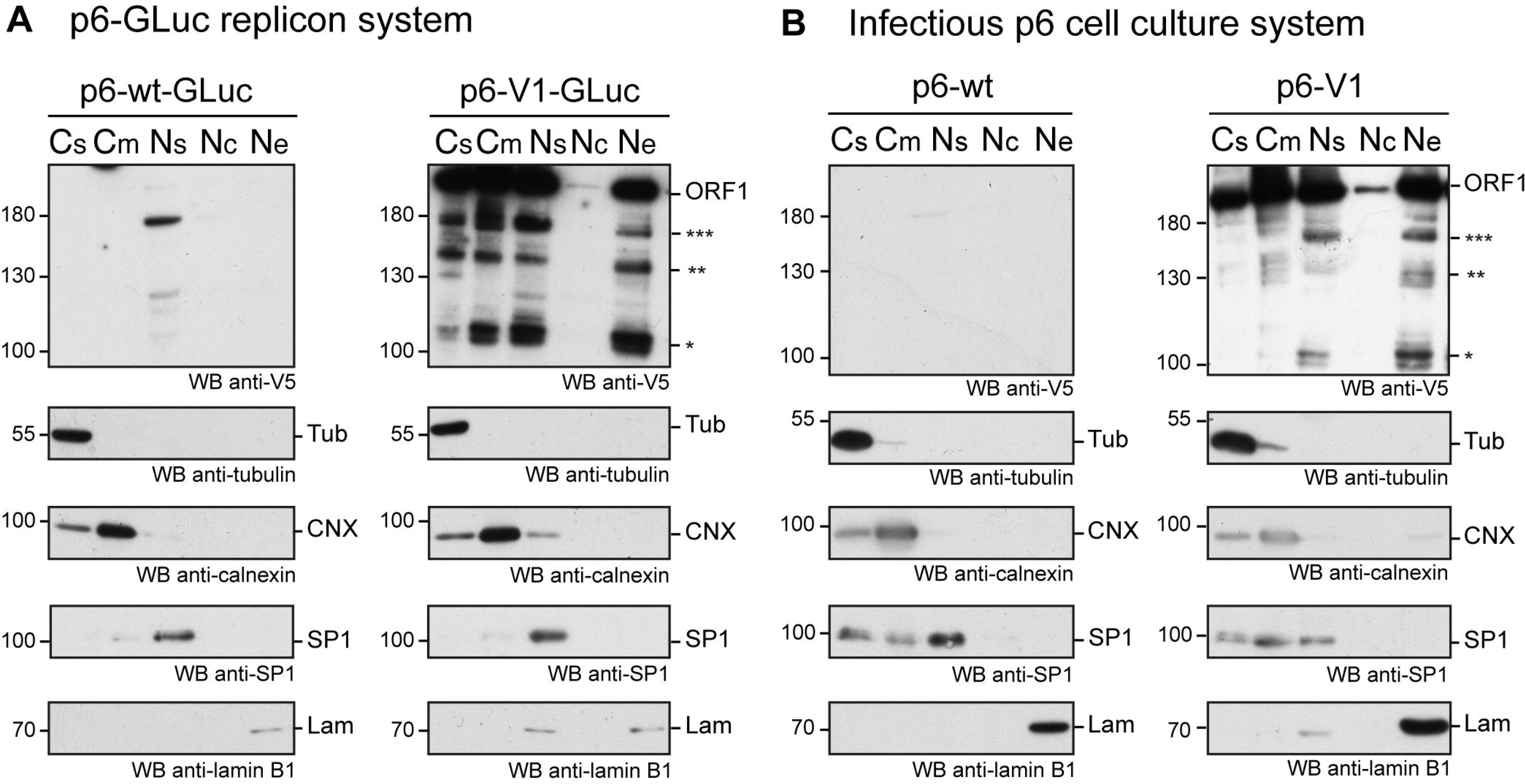
The ORF1 protein is expressed in different cellular compartments. **Figure 5A.** PLC3 cells were electroporated with the p6-GLuc replicons expressing the non-tagged wildtype ORF1 (p6-wt-GLuc), and the V5-tagged ORF1 (p6-V1-GLuc). Three dpe, cells were collected and cellular fractions were separated as follows: Cs = cytoplasmic soluble fraction, Cm = cytoplasmic membranous fraction, Ns = nuclear soluble fraction, Nc = nuclear chromatin-bound fraction, Ne = Nuclear envelope. Immunoblots were probed with a mouse monoclonal antibody directed against the V5 epitope. Other antibodies were used to monitor for fraction enrichment: anti-γ-tubulin as a marker of the cytoplasmic soluble fraction, anti-calnexin as a marker of cytoplasmic membranous fraction, anti-SP1 as a marker of nuclear soluble fraction and anti-lamin B1 as a marker of nuclear envelope. Molecular weight markers are indicated in kilodaltons. Stars indicate lower molecular weight ORF1 products. **Figure 5B.** PLC3 cells were electroporated with the p6 expressing the non-tagged wildtype ORF1 (p6-wt), and the V5-tagged ORF1 (p6-V1). Four hpe, cells were collected and cellular fractions were separated and named as above. The above-mentioned antibodies were used to reveal the immunoblots. Molecular weight markers are indicated in kilodaltons. Stars indicate lower molecular weight ORF1 products.

### V5-tagged HEV p6 remains infectious in cell culture

We also determined the impact of V5 insertions on HEV infectivity. Huh-7.5 cells were infected with the supernatant of PLC3 cells that were electroporated with p6-wt, p6-V1 or p6-V2 RNAs (**Figure 6**). Three days post-infection, Huh-7.5 cells were processed for anti-ORF2 indirect immunofluorescence. ORF2-positive cells were counted and each positive cell focus was considered as one focus forming unit (FFU). When compared to the p6-wt, p6-V1 and p6-V2 strains produced infectious titers that were not significantly different (**Figure 6A**, respective Mann-Whitney pvalues of 1 and 0.4), indicating that V5-epitope insertions did not alter HEV assembly and infectivity. Analysis by confocal microscopy also showed similar numbers of ORF2-positive cells and staining patterns when comparing Huh-7.5 cells that were infected with p6-wt *versus* V5-tagged p6 (**Figure 6B**).

**Figure 6.**
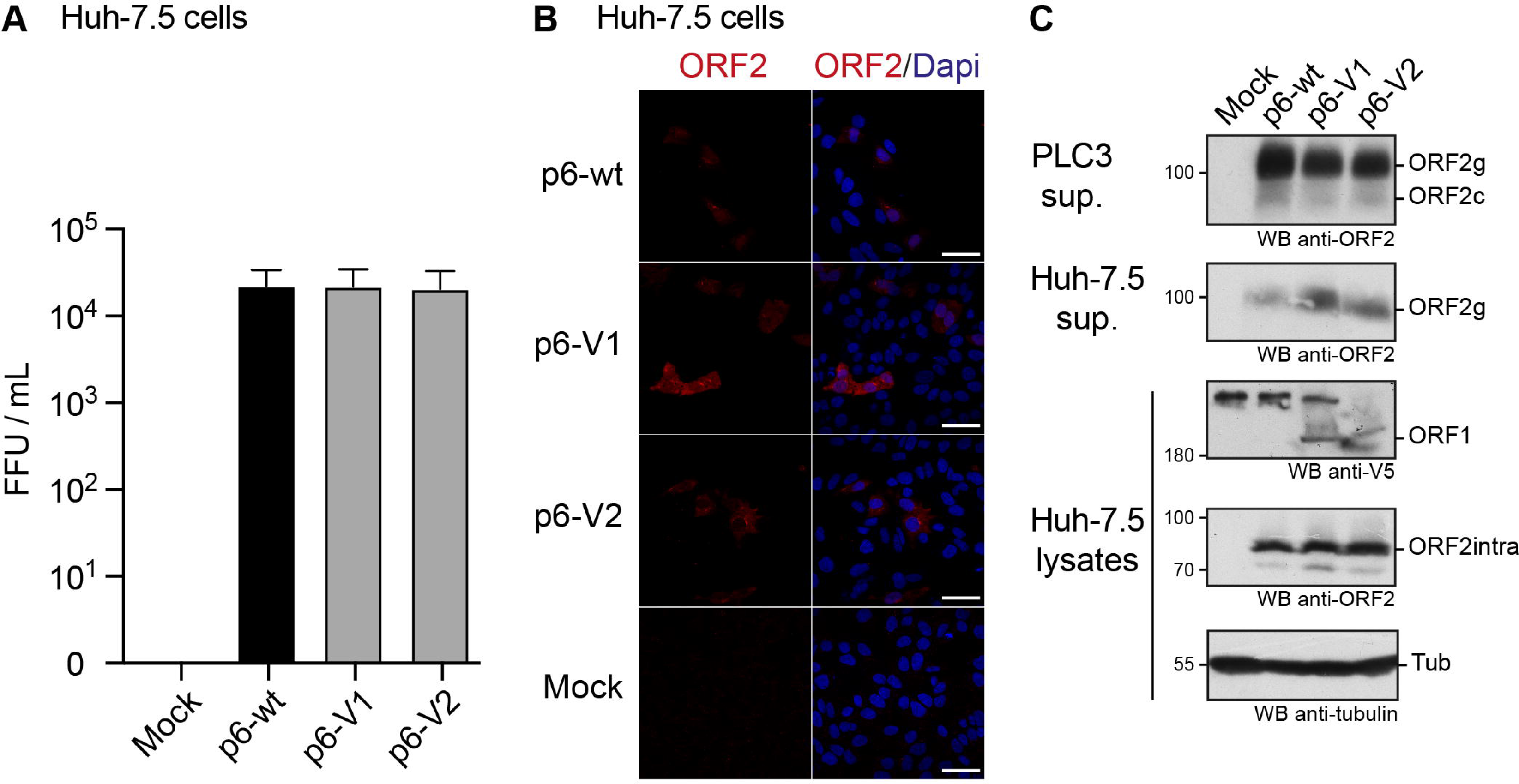
Infection of Huh-7.5 cells by HEV p6 expressing V5-tagged ORF1 polyprotein. **Figure 6A.** Infectious titers of HEV p6 expressing epitope-tagged ORF1 polyprotein. Ten dpe, supernatants of PLC3 cells electroporated with p6-wt, p6-V1 or p6-V2 were collected to infect Huh-7.5 cells. Three days post-infection, Huh-7.5 cells were fixed and stained with an anti-ORF2 antibody (1E6). The number of ORF2-positive cells from 3 independent experiments were counted and used to calculate infectious titers in focus forming unit (FFU/mL). Statistical analysis was performed using the Mann-Whitney test. **Figure 6B.** Immunofluorescence of HEV p6-infected Huh-7.5 cells. Huh-7.5 cells were infected with the supernatants of PLC3 cells that were electroporated with HEV p6 expressing non-tagged ORF1 (p6-wt) or V5-tagged (p6-V1, p6-V2) replicase as mentioned above. Three days post-infection, cells were processed for anti-ORF2 immunofluorescence (1E6, red) prior to analysis by confocal microscopy. Cell nuclei were stained with Dapi (blue). Huh-7.5 cells infected with mock-electroporated PLC3 cell supernatant served as negative control (Mock). Scale bar = 50 μm. **Figure 6C.** Expression of epitope-tagged ORF1 and ORF2 proteins in the p6 cell culture system. PLC3 cells were electroporated with p6-wt, p6-V1 or p6-V2. Ten dpe, total protein extracts were submitted to Western blot and probed with an anti-ORF2 antibody (1E6, first panel, PLC3 sup.). Huh-7.5 cells were infected with PLC3 supernatants. Three days post-infection, Huh-7.5 supernatants were also analyzed for ORF2 expression (1E6, second panel, Huh-7.5 sup.). In parallel, Huh-7.5 cells were lysed and total protein extracts were probed with anti-V5, -ORF2 (1E6) and -γ-tubulin antibodies (last 3 panels, Huh-7.5 lysates). Molecular weight markers are indicated in kilodaltons. ORF1 = full-length protein, ORF2g = glycosylated form of ORF2, ORF2c = cleaved form of ORF2, ORF2intra = intracellular ORF2 form, Tub = γ-tubulin.

Of note, at 10 dpe, ORF2 protein expression was also controlled by immunoblot in the PLC3 cell supernatants that were used to infect Huh-7.5 cells (**Figure 6C**, PLC3 sup.). The ORF2 protein was detected as efficiently in all PLC3 supernatants irrespective of the electroporated p6 strain. After 3 days of infection with PLC3 supernatants producing p6-wt and V5-tagged-p6 particles, the ORF2 protein was also detected in lysates and supernatants of Huh-7.5 cells (**Figure 6C**, Huh-7.5 sup., Huh-7.5 lysates). Moreover, the V5-tagged ORF1 protein could also be detected in Huh-7.5 cell lysates. These data confirm that the V5-epitope insertions do not affect production and infectivity of HEV particles.

### Subcellular localization of the epitope-tagged ORF1 replicase and co-localization with ORF2 and ORF3 proteins

We next took advantage of replicative epitope-tagged ORF1 constructs to analyze the subcellular localization of ORF1 replicase by confocal imaging. PLC3 cells were electroporated with p6-wt, p6-V1 and p6-V2 RNAs. At 3 dpe, cells were first processed for single V5 staining. The V5 antibody mostly displayed a nuclear staining as well as cytoplasmic accumulation in the vicinity of the nucleus (**Figure 7A**, white arrowheads). Secondly, double immunostainings with antibodies directed against the V5 epitope and ORF2 or ORF3 proteins were performed. Interestingly, the ORF2 and ORF3 stainings partially overlapped with the V5 staining in peri-nuclear nugget-like substructures (**Figures 7C,E**), a finding that was corroborated by calculating the Pearson’s correlation coefficients (PCC, **Figure 7B**). In line with these observations, super-resolution confocal microscopy analyses of PLC3 cells electroporated with p6-V1 and co-labeled with either V5 / ORF2 or V5 / ORF3 antibodies, showed a total overlap of fluorescence intensities between the V5 signal and both the ORF2 and ORF3 signals (**Figures 7D,F**). Altogether these results indicate that V5-tagged ORF1, ORF2 and ORF3 proteins are co-distributed in compact structures located in the vicinity of the nucleus of HEV-producing cells.

**Figure 7.**
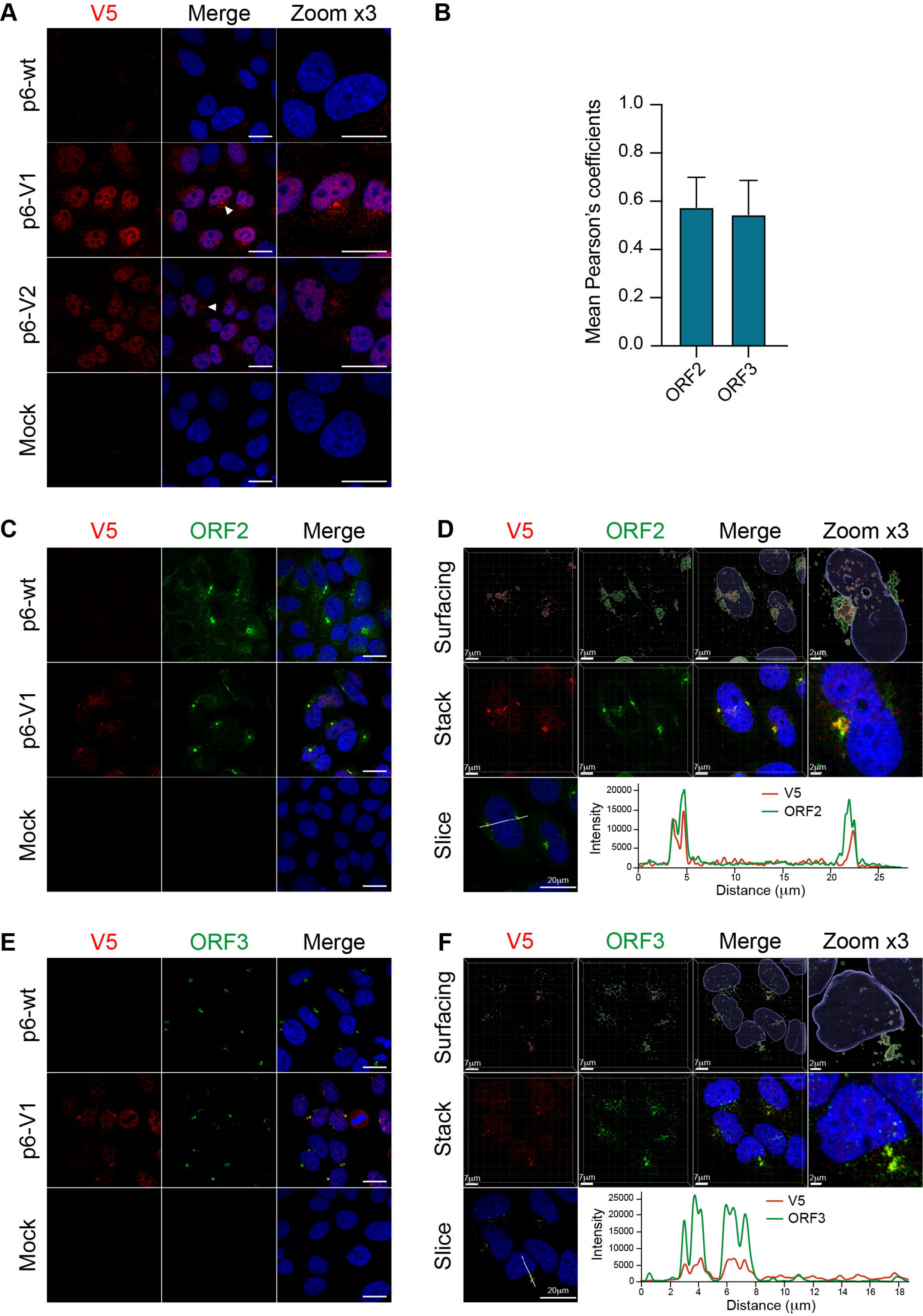
Co-localization of the V5-tagged ORF1 and ORF2/ORF3 proteins in the host cell. **Figure 7A.** PLC3 cells were electroporated with the p6 strain expressing either the untagged (p6-wt) or the V5-tagged ORF1 (p6-V1, p6-V2) proteins. Three dpe, cells were processed for immunofluorescence using an anti-V5 antibody (red) prior to analysis by confocal microscopy. Perinuclear nugget-like structures are shown (white arrowheads). The cell nuclei were stained with DAPI (blue). Mock-electroporated cells served as negative control. Scale bar = 20 μm. **Figure 7B.** Regions of interest (ROI) were drawn around the perinuclear nugget-like structures on images taken from PLC3 cells electroporated with p6-V1 and co-labeled with antibodies directed against the V5 epitope and the ORF2 (**Figure 7C**) or the ORF3 proteins (**Figure 7E**). ROI were used to determine Pearson’s correlation coefficients (PCC) between V5-tagged ORF1 and ORF2 (ORF2) labeling or V5-tagged ORF1 and ORF3 (ORF3) labeling using JACoP plugin from ImageJ software. PCC means (± standard deviation) were calculated from 30 different ROI. **Figures 7C-7E.** Co-labeling of the V5-tagged HEV ORF1 replicase with ORF2/ORF3 in PLC3 cells. PLC3 cells were electroporated with p6-wt or p6-V1. Three dpe, viral proteins were co-labeled with antibodies directed against the V5 epitope (red, V5) and (i) ORF2 (1E6, green, **Figure 7C**) or (ii) ORF3 (green, **Figure 7E**) prior to analysis by confocal microscopy. Mock-electroporated cells served as negative control. Antibodies used are listed in **Table 2**. Scale bar = 20 μm. **Figures 7D-7F.** PLC3 cells electroporated with p6-V1 and co-stained with anti-V5 and (i) anti-ORF2 (**Figure 7D**) or (ii) anti-ORF3 antibodies (**Figure 7F**) were analyzed by confocal microscopy with a high resolution Airyscan module. On the top, volume rendering of the 3D z-stacks (Surfacing) using the Imaris software are shown to visualize the V5-tagged ORF1/ORF2 (**Figure 7D**) or ORF1/ORF3 (**Figure 7F**) substructures. In the middle, z-stacks are shown. On the bottom, line graphs show the fluorescence intensities of V5-tagged ORF1 and ORF2 or ORF3 staining measured every 50 nm across the region of interest highlighted by the white line in the micrograph shown on the left. Scale bars show the indicated length.

**Figure 8.**
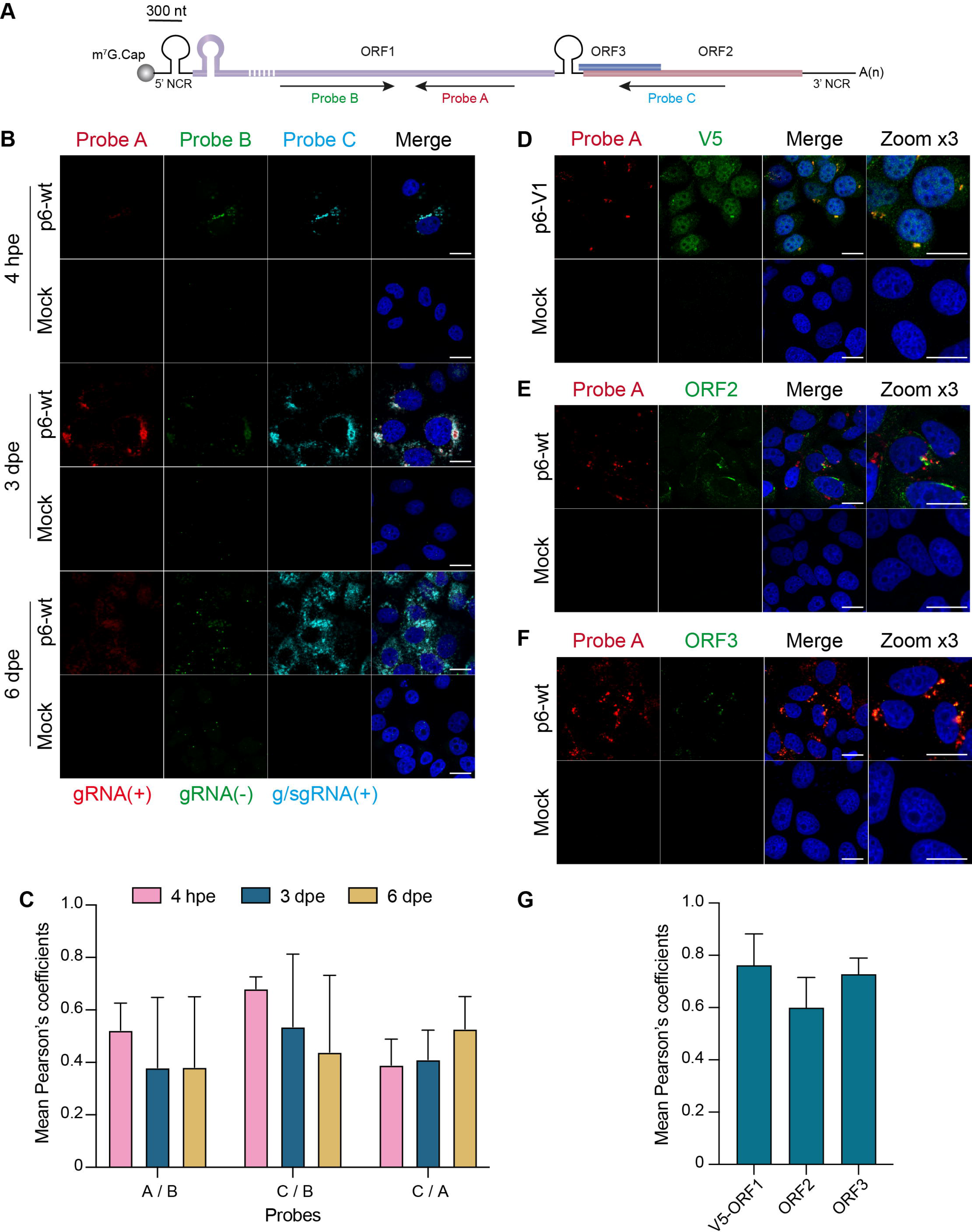
*In situ* labeling of positive- and negative-sense HEV RNAs. PLC3 cells were electroporated with untagged (p6-wt) or V5-tagged (p6-V1) p6 strains. Cells were grown on coverslips, fixed at 4 hpe, 3 and 6 dpe and processed for *in situ* RNAscope® hybridization. Cell nuclei were stained with Dapi (blue). Images were taken on a confocal microscope. Mock-electroporated cells served as negative control. Scale bar = 20 μm. **Figure 8A.** Schematic overview of the RNAscope® probe location. Probe A targets the positive-sense genomic RNA and is located in the RdRp domain of ORF1 (purple). Probe B targets the negative-sense RNA and is also located in the RdRp but does not overlap with probe A. Probe C targets the positive-sense subgenomic RNA, by hybridizing at the ORF3 (blue) / ORF2 (red) overlap. The full-length of the ORF1 protein cannot be represented at the scheme scale (purple dashed lines). **Figure 8B.** PLC3 cells electroporated with the p6-wt strain were sequentially stained with probes A (red, gRNA(+)), B (green, gRNA(-)) and C (cyan, g/sgRNA(+)) at 4 hpe, 3 and 6 dpe. gRNA(+) = positive-stranded genomic RNA; gRNA(-) = negative-stranded genomic RNA; g/sgRNA(+) = positive-stranded genomic and subgenomic RNAs. **Figure 8C.** Immunofluorescence images of whole PLC3 cells electroporated with p6-wt and co-labeled with probes A, B and C were used to determine PCC using JACoP plugin from ImageJ software. PCC means (± standard deviation) were calculated from 30 analyzed whole cells. **Figures 8D-8F.** Three dpe, PLC3 cells electroporated with the p6-V1 construct were stained with probe A (red) and (i) the anti-V5 antibody (green, **Figure 8D**), (ii) the anti-ORF2 (green, 1E6, **Figure 8E**) or (iii) the anti-ORF3 (green, **Figure 8F**). **Figure 8G.** Regions of interest (ROI) were drawn around the perinuclear nugget-like structures on images taken from PLC3 cells electroporated with p6-V1 or p6-wt and co-labeled with probe A and anti-V5, anti-ORF2 or anti-ORF3 antibody. Twenty ROI were used to calculate mean PCC (± standard deviation) between probe A and V5-tagged ORF1 (V5-ORF1) labeling or probe A and ORF2 labeling (ORF2) or probe A and ORF3 labeling (ORF3) using JACoP plugin from ImageJ software.

Since the ORF1 protein has been largely studied in heterologous expression systems, we next wanted to compare the subcellular localization of V5-tagged ORF1 in replicative and heterologous systems. Eight hours post-transfection with the V5-tagged constructs, a mostly cytoplasmic reticular labeling was observed in H7-T7-IZ cells (**Supplementary Figure 3A**). Also, in H7-T7-IZ cells co-transfected with the V5-tagged ORF1 and ORF2/ORF3 expressing plasmids, partial staining overlaps of the V5-tagged replicase with ORF2 and ORF3 proteins were visible in the cytoplasm but also in perinuclear accumulations (**Supplementary Figures 3B,C**). Thus, the heterologous system recapitulates the partial co-distribution of V5-tagged ORF1 with ORF2/ORF3 that was also observed in perinuclear substructures of HEV-producing cells.

In order to locate HEV replication sites, we implemented a highly sensitive technology (RNAscope®) to detect viral RNA. Three probes were designed to specifically hybridize to the positive- and negative-HEV RNA strands (**Figure 8A**). The probes A and C target the positive strand of the genomic and subgenomic HEV RNAs, respectively. The probe B targets the negative strand of the genomic RNA. The p6-wt-electroporated PLC3 cells were fixed at 4 hpe, 3 and 6 dpe and, then submitted to *in situ* hybridization using probes A, B and C sequentially. A fluorescent signal was detected for each of the 3 probes in discrete perinuclear foci in the host cell as early as 4 hpe (**Figure 8B**). While fluorescent staining of the negative-sense RNA (probe B) and subgenomic RNA (probe C) overlapped (PCC_B/C_ = 0.68 ± 0.05, **Figure 8C**), the staining of probe A only partially overlapped with the formers (PCC_B/A_ = 0.52 ± 0.10, PCC_A/C_ = 0.39 ± 0.10, **Figure 8C**). Three dpe, the positive-stranded-RNA staining (probes A and C) further expanded all around the cell nuclei while the negative-stranded-RNA staining (probe B) remained more condensed as perinuclear foci. At 6 dpe, the staining of genomic and subgenomic positive-sense RNAs adopted a diffuse pattern in the cytoplasm with a PCC_A/C_ reaching 0.53 ± 0.12. The negative-stranded-RNA staining was closer to a dot-like fainter pattern while the staining overlap with the 2 other probes decreased (PCC_A/B_ = 0.38 ± 0.27, PCC_B/C_ = 0.44 ± 0.29). At every time points, the probe C staining surrounded the probe A staining while the probe B staining was the faintest of all.

Next, positive-sense genomic RNA (probe A) were co-labeled with the viral proteins ORF1, ORF2 and ORF3 (**Figures 8D-F**). At 3 dpe, the V5-tagged ORF1 protein co-localized with the positive-sense genomic RNA within the previously identified perinuclear foci with a strong PCC of 0.76 ± 0.11 (**Figures 8D,G**). The viral RNA also co-distributed with ORF2 and ORF3 proteins at 3 dpe in p6-wt-electroporated PLC3 cells (**Figure 8G**, PCC_A/ORF2_ = 0.60 ± 0.12, PCC_A/ORF3_ = 0.73 ± 0.06). These accumulations were also located in the perinuclear proximity of the cell (**Figures 8E,F**).

### Identification of the HEV-induced substructures

Recently, the ORF2 and ORF3 proteins were reported to colocalize with cellular markers of the endocytic recycling compartment (ERC) in perinuclear substructures (Bentaleb et al., 2021). To further delineate whether the V5-tagged ORF1 was also present in these substructures, we conducted co-labeling experiments of the V5-tagged-ORF1 with ERC markers such as CD71, Rab11, EHD1 and PACSIN2 in PLC3 cells electroporated with p6-V1 (**Figure 9, Supplementary Figure 3D**). The V5-tagged ORF1 staining overlapped with CD71 and Rab11 staining in the perinuclear substructures (**Figures 9A,C**). Analyses of super-resolution confocal microscopy images further strengthened these observations showing a total overlap of fluorescence intensities between V5-tagged-ORF1 and CD71 (**Figure 9B**) as well as between V5-tagged ORF1 and Rab11 (**Figure 9D**) in the perinuclear substructures. The V5-tagged ORF1 colocalized best with Rab11 (PCC = 0.67 ± 0.08) whereas the colocalization was moderate with CD71 and EHD1 (PCC = 0.42 ± 0.13 and 0.49 ± 0.10, respectively, **Figure 9F**). No colocalization was found between V5-tagged ORF1 and PACSIN2 (PCC = 0.22 ± 0.11, **Figure 9F**, **Supplementary Figure 3D**). In addition, contrary to a previous report (Szkolnicka et al., 2019; Bentaleb et al., 2021), no colocalization was found between V5-tagged ORF1 and CD63 or CD81, two markers of multivesicular bodies, while CD81 labeling appeared surrounding the V5-tagged ORF1 signal (**Figure 9F**, **Supplementary Figures 3E-F)**.

**Figure 9.**
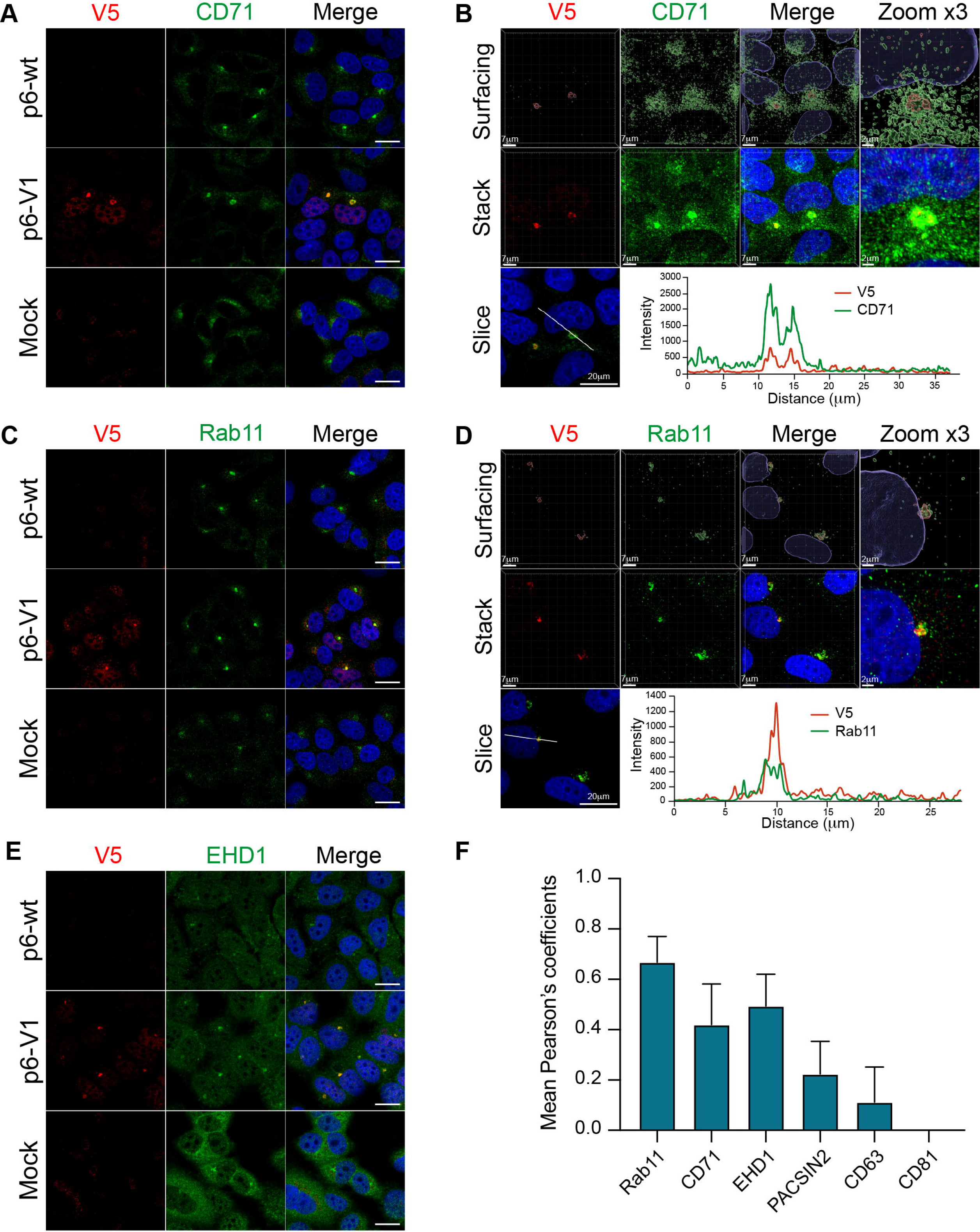
Co-labeling of the HEV V5-tagged ORF1 protein with several cellular markers. The wt (p6-wt) and V5-tagged (p6-V1) ORF1 proteins were expressed in PLC3 cells electroporated with the p6 HEV strain. Three dpe, cells were co-labeled with anti-V5 antibody (red) and cellular markers antibodies (green) directed against CD71 (**Figures 9A,B**), Rab11 (**Figures 9C,D**), EHD1 (**Figure 9E**). Cell nuclei were stained with DAPI (blue). Mock-electroporated cells served as negative controls (Mock). Images were taken with a confocal microscope. Antibodies used are listed in **Table 2**. Scale bar = 20 μm. PLC3 cells electroporated with p6-V1 and co-stained with anti-V5 and antibodies directed against CD71 (**Figure 9B**) and Rab11 (**Figure 9D**) were analyzed by confocal microscopy with a high resolution Airyscan module. On the top, volume rendering of the 3D z-stacks (Surfacing) using the Imaris software are shown to visualize the V5-tagged ORF1/CD71 or Rab11 substructures. In the middle, z-stacks are shown. On the bottom, line graphs show the fluorescence intensities of V5-tagged ORF1 and CD71/Rab11 staining measured every 50 nm across the region of interest highlighted by the white line in the micrograph shown on the left. Scale bars show the indicated length. **Figure 9F**: Regions of interest (ROI) were drawn around the perinuclear nugget-like structures on the immunofluorescence images of p6-V1 electroporated PLC3 cells co-labeled with antibodies directed against the V5 epitope and CD71, Rab11, EHD1 (**Figures 9A,C,E**) and PACSIN2, CD63 and CD81 (**Supplementary Figures 3D-F**). Thirty ROI were used to calculate PCC (+ standard deviation) using JACoP plugin from ImageJ software.

## Discussion

Due to its low expression level, its tight regulation in time and space as well as a lack of a commercial antibody (Lenggenhager et al., 2017), the non-structural ORF1 protein is the least studied of the three HEV proteins. Thus, the insertion of epitope tags within the HEV replicase appeared as a practical strategy to characterize the subcellular localization and processing of ORF1 polyprotein. Indeed, a transposon-based approach to insert HA epitopes in the ORF1 genome has been recently used to characterize the subcellular localization of the HEV replicase (Szkolnicka et al., 2019). Similarly, we aimed at finding positions within ORF1 where epitope tags could be inserted without disturbing viral replication. After aligning 44 HEV strains, the HVR appeared as the least conserved domain in the ORF1 sequence. In addition, while presenting the highest divergence of the entire HEV genome (Muñoz-Chimeno et al., 2020), the HVR is known to tolerate inserted fragments arising either from duplication of viral genome or from human genes which were reported to confer better replication efficacies or adaptation to cell culture (Shukla et al., 2011, 2012; Nguyen et al., 2012; Johne et al., 2014; Lhomme et al., 2014).

Recently, a cell culture model derived from the PLC/PRF/5 cell line has been established in the laboratory to efficiently produce HEV particles from the Kernow C-1 p6 strain (Shukla et al., 2012; Montpellier et al., 2018). In a first approach to identify non-disruptive insertion sites, the ORF1 HVR aa sequence of the Kernow C-1 p6 strain was aligned with those of common epitope tags. Four aa of the HVR were matching with the V5 tag sequence, while 3 aa had to be mutated and 7 had to be inserted to construct the full V5 epitope aa sequence (V1, **Figure 1**). Secondly, a V5 or HA epitope (V2 and H2, **Figure 1**) were inserted into the Kernow C-1 p6 strain in the position where the S19 insertion was found in a different strain (LBPR-0379) to confer a replication advantage in cell culture (Nguyen et al., 2012; Shukla et al., 2012). Lastly, an HA epitope was inserted at the C-terminus of the ORF1 coding sequence to avoid impacting the structure of the ORF1 protein and thereby perturb replication (H1, **Figure 1).**

At first, the epitopes were inserted into the p6-GLuc replicon in order to determine the impact of the tag insertion on its replicative ability. The luciferase activity was measured over time. Result analyses led to the conclusion that the H1 insertion diminishes the expression of the subgenomic genes, as luciferase activity was greatly decreased, which is likely due to the disruption of the subgenomic promoter region (Ding et al., 2018). Therefore, insertion at the H1 position is likely to also impact replication in the p6 context but could be of use in the heterologous expression system. Next, H2 and V2 insertions, which were placed in the same position, impacted the subgenomic expression differently. While the p6-V2-GLuc showed similar luciferase activity to the non-tagged p6-wt-GLuc, the p6-H2-GLuc construct showed a decreased replication efficacy. Thus, the aa sequence composition of the epitope, and not only the insertion position, may impact the replicase activity by modifying its conformational structure. Out of the 4 tagged replicons tested, replication efficacies of the p6-V1-GLuc and p6-V2-GLuc constructs appeared the least affected by epitope insertion. The luciferase activity of these constructs resembled those of the non-tagged p6-wt-GLuc. Moreover, the quantification of extracellular and intracellular viral RNA over the course of 7 days displayed similar kinetics in PLC3 cells expressing the p6-wt or the V5-tagged p6 constructs, thus strengthening the fact that the V5 insertions within the ORF1 HVR did not disturb the HEV replication. In line with these results, the infectivity of the V5-tagged p6 constructs was not altered when compared to p6-wt. Therefore, the V5-tagged constructs were selected to delineate the ORF1 features.

One of our goal was to characterize the processing of the ORF1 protein. Polyproteins encoded by positive stranded RNA viruses are commonly subjected to cleavage by viral and/or cellular proteases (Ploss and Dubuisson, 2012; Gu and Rice, 2013; Baggen et al., 2021; V’kovski et al., 2021). However, in the HEV field, the literature on ORF1 cleavage is highly controversial as there is evidence for and against cleavage of the non-structural protein (LeDesma et al., 2019).

In an aim to study potential processing of HEV replicase, expression of the V5-tagged ORF1 protein was analyzed over time by immunoblotting. In the 3 different systems tested in this study, the full-length ORF1 protein and smaller bands that may correspond to potential ORF1 cleavage products were detected, especially at earlier time points. This could reflect the early need for ORF1 to replicate RNA at the onset the viral cycle. Although full-length ORF1 protein was identified by mass spectrometry with certainty, full sequence identification of the smaller products was less robust due to their lower expression levels. The ORF1 protein expressed heterologously does not seem to be processed from its N-terminus while in the p6 infectious system, more ORF1 potential cleavage products were detected and the N-terminus of the protein also seemed more stable than the C-terminus. The expression of more numerous potential cleavage products in the p6 infectious system may suggest the requirement of all viral proteins as well as cellular proteins to achieve the full ORF1 processing. As the possibility of degradation cannot be fully excluded, we inhibited the proteasome with lactacystin and noted that the observed band pattern and ORF1 quantity remained unchanged. Thus, the minor ORF1 products may be the result of natural viral and/or cellular processing rather than artefactual degradation. Furthermore, the ORF1 protein and its potential cleavage products were detected in different cellular compartments with slightly different patterns. For example, a band of 160 kDa was only present in the nuclear fraction enriched from the p6 cell culture system. Nevertheless, without any sequence information, it remained difficult to assess to which domain of the ORF1 protein this product may correspond.

RNA viruses generally replicate in the cytoplasm. In previous studies, the ORF1 protein was found in the cytoplasm colocalizing with ORF2 and ORF3 viral proteins, and ERGIC/Golgi markers (Rehman et al., 2008; Szkolnicka et al., 2019). In our study, the V5-tagged ORF1 proteins, expressed in the replicon, heterologous and p6 cell culture expression systems, displayed both a cytoplasmic and nuclear localization as assessed by subcellular fractionation and immunofluorescence experiments. The use of the Kernow C-1 p6 strain, that contains the S17 human ribosomal protein insertion in which an element could act as a nuclear localization signal (NLS), may account for this discrepancy (Kenney and Meng, 2015). However, mutations of the conserved aa in the S17 NLS disrupted the replication efficacy of the p6-GLuc replicon and did not inhibit the nuclear localization as assessed by immunofluorescence staining or immunoblot analysis of subcellular fractions in the heterologous system (data not shown). In addition, we observed perinuclear aggregations of the ORF1 protein that often coincide with a deformation of the nucleus. Recently, ORF2 and ORF3 proteins as well as cellular markers of the ERC such as Rab11 and CD71 were reported to locate to this region in HEV-producing cells (Bentaleb et al., 2021). Interestingly, we confirmed partial overlaps of the ORF1 protein with ORF2 and ORF3 viral proteins as well as several cellular markers of the ERC, especially in the nugget-like perinuclear region. These results indicate that the place of viral replication is in close proximity to the site of virus assembly. Indeed, in other viruses such as Dengue virus, replication and assembly take place in the same subcellular compartment (Welsch et al., 2009).

Lastly, to formerly identify the HEV replication site, we aimed at locating the negative-sense HEV RNA in the host cell. To that end, the RNAscope® technique, which enabled to specifically target and visualize the positive- and negative-RNA-strands of hepatitis C virus and the positive-RNA-strand of Zika virus, was implemented (Wang et al., 2012; Liu et al., 2019). We managed to locate positive-sense genomic and subgenomic HEV RNAs as well as the negative-sense RNA in the cell. Moreover, co-distributions of the positive-sense genomic RNA with ORF1, ORF2 and ORF3 viral proteins were visible within the nugget-like perinuclear foci. In conclusion, we demonstrated that viral replication and assembly take place in close proximity in HEV-producing cells, in perinuclear nugget-like structures that may constitute the HEV viral factories.

### Author Contributions

K.M., C.B., K.H., C.M., V.A., C.M., J-M.-F., M.F., C.C., Y.R., C.L., J.D., L.C. and C-M.A-D. performed research and/or analyzed data.

K.M. and C-M.A-D. wrote the paper.

## Acknowledgments

We thank Suzanne U. Emerson (NIH, USA), Jérôme Gouttenoire (University of Lausanne) and Ralph Bartenschlager (University of Heidelberg) for providing us with reagents. We thank the imaging core facility of the BioImaging Center Lille-Nord de France for access to the instruments.

