## Supplementary Information for "Characterization of the hepatitis E virus replicase"

### SUPPLEMENTARY MATERIAL

**Supplementary Figure 1**. Replication inhibition of p6-GLuc replicons and p6 infectious strains in PLC3 and Huh-7.5 cells in presence of sofosbuvir.

**Supplementary Figures 1A,B.** Replication efficacies of HEV replicons expressing tagged ORF1 in PLC3 and Huh-7.5 cells treated with sofosbuvir. Cells were electroporated with the tagged p6-GLuc replicons (V1, V2, H1, H2) or untagged p6-GLuc replicons (wt). Four hpe, sofosbuvir, an effective polymerase inhibitor, was added to the cell medium at a final concentration of 20 µM. The relative light units (RLU) were measured at 5 dpe by quantification of the *Gaussia* luciferase in the cell supernatants using a luminometer. Replication fold increases were normalized to 1 dpe. Percentages of replication were calculated to the non-treated corresponding construct. Experiments were conducted three times with 3 technical replicates.

**Supplementary Figure 1C**. Immunofluorescence of p6-electroporated PLC3 cells in presence of sofosbuvir (20 µM). Three dpe with the infectious p6-wt, p6-V1 or p6-V2 constructs, PLC3 cells were processed for immunostaining using an anti-ORF2 antibody (red, 1E6, **Table 2**) prior to confocal microscopy analysis. Cell nuclei were stained with Dapi (blue). Mock-electroporated PLC3 cells served as negative control. Scale bar = 50 µM.

**Supplementary Figure 2. Proteasome inhibition by lactacystin.**

H7-T7-IZ cells were transfected with the pTM expressing the untagged ORF1 (ORF1-wt) or the V5-tagged ORF1 (ORF1-V1, **Supplementary Figure 2A**). PLC3 cells were electroporated with the p6-GLuc replicons expressing the untagged ORF1 (p6-wt-GLuc) or the V5-tagged ORF1 (p6-V1-GLuc, **Supplementary Figure 2B**). One dpt of H7-T7-IZ and 3 dpe of PLC3 cells, a specific proteasome inhibitor lactacystin (30 μM, Sigma Aldrich) was applied for 8 hours and total cell lysates were collected in presence of protease inhibitors. The band migrating higher than 180 kDa corresponds to the full-length ORF1 protein (ORF1). Mock-transfected cells served as negative control (Mock). Immunoblots were probed with antibodies directed against the V5 epitope, HSP70 and γ-tubulin (Tub) proteins (**Table 2**). HSP70 and γ-tubulin are detected to monitor for efficient proteasome inhibition and for even protein loading, respectively. NT: non-treated samples; V: vehicle-treated samples (methanol); Lact: lactacystin-treated samples. Molecular weight markers are indicated in kilodaltons. Stars indicate lower molecular weight ORF1 products.

**Supplementary Figure 3. Co-localization of the HEV proteins in the host cell.**

**Figure 3A**. Eight hpt, H7-T7-IZ cells expressing either the wildtype ORF1 (ORF1-wt) or the V5-tagged ORF1 proteins (ORF1-V1, ORF1-V2) were processed for anti-V5 immunostaining (red, **Table 2**) prior to analysis by confocal microscopy. Cell nuclei were stained with DAPI (blue). Mock-transfected cells served as negative control. Scale bar = 20 µm.

**Figures 3B,C**. Co-labeling of the V5-tagged HEV replicase with ORF2 or ORF3 in H7-T7-IZ cells. Cells were treated as above. Then, the V5-tagged ORF1 protein (V5 antibody, red) was co-labeled with the ORF2 (green, **Figure 3B**) or the ORF3 (green, **Figure 3C**) proteins prior to analysis by confocal microscopy. Used antibodies are listed in **Table 2**. Scale bar = 20 µm.

**Figures 3D-F.** Co-labeling of the HEV V5-tagged ORF1 protein with several cellular markers. The wt (p6-wt) and V5-tagged (p6-V1) ORF1 proteins were expressed in PLC3 cells electroporated with the p6 HEV infectious strain. Three dpe, cells were co-labeled with anti-V5 antibody (red) and cellular markers antibodies (green) directed against PACSIN2 (**Figure 3D**), CD63 (**Figure 3E**) and CD81 (**Figure 3F**) prior to confocal microscopy analysis. Cell nuclei were stained with DAPI (blue). Mock-electroporated cells served as negative controls (Mock). Scale bar = 20 µm.
