## Supplementary figures and images for "Characterization of the hepatitis E virus replicase"

### Supplementary Figure 1

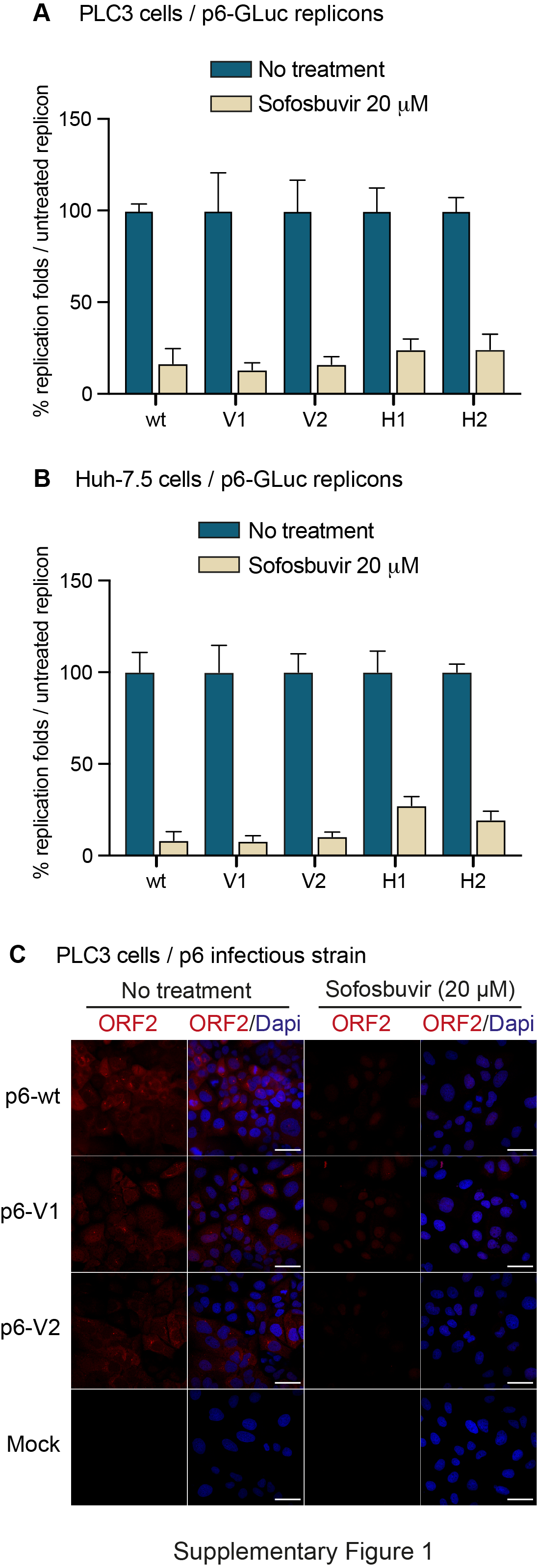

### Supplementary Figure 2

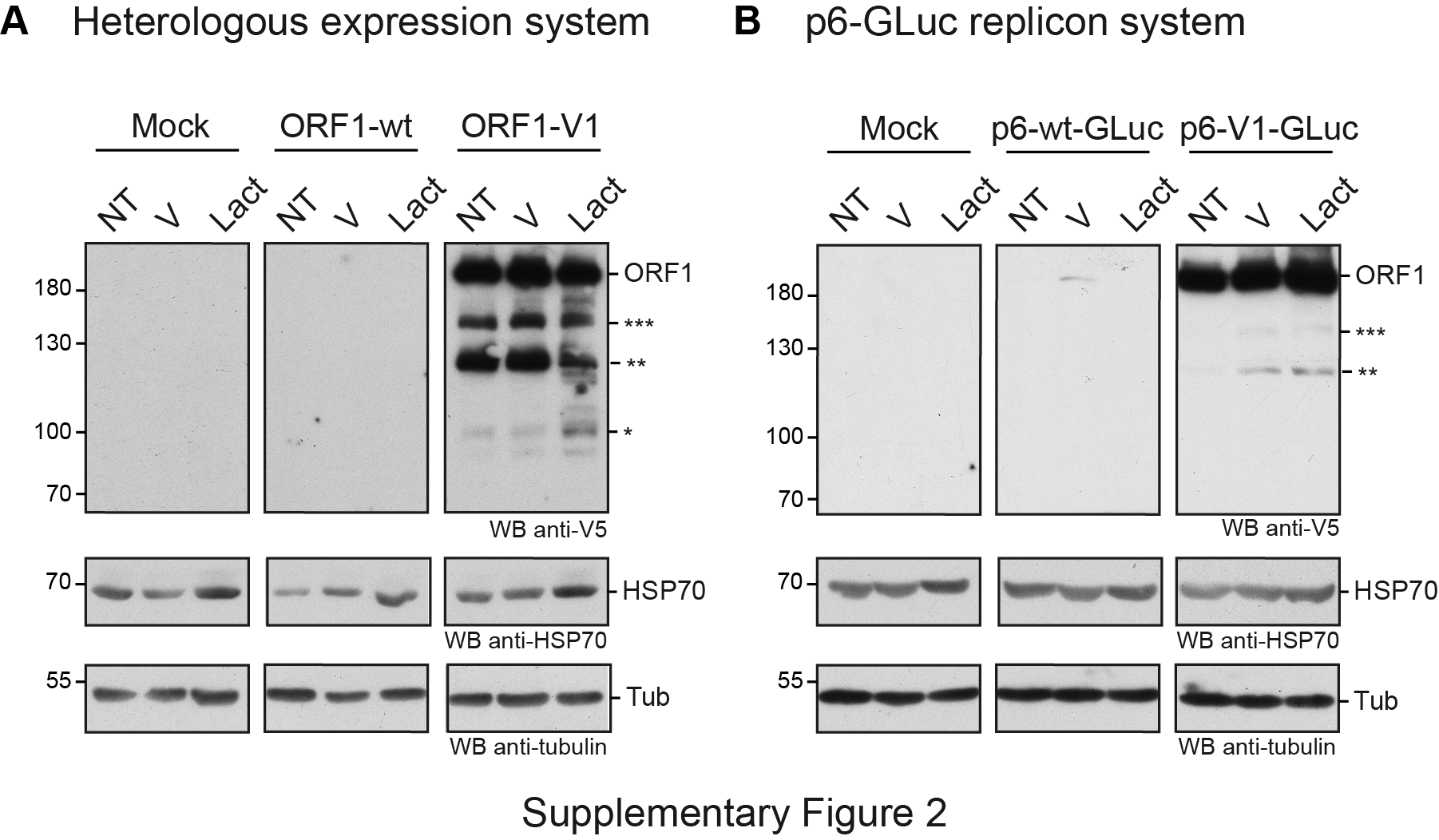

### Supplementary Figure 3

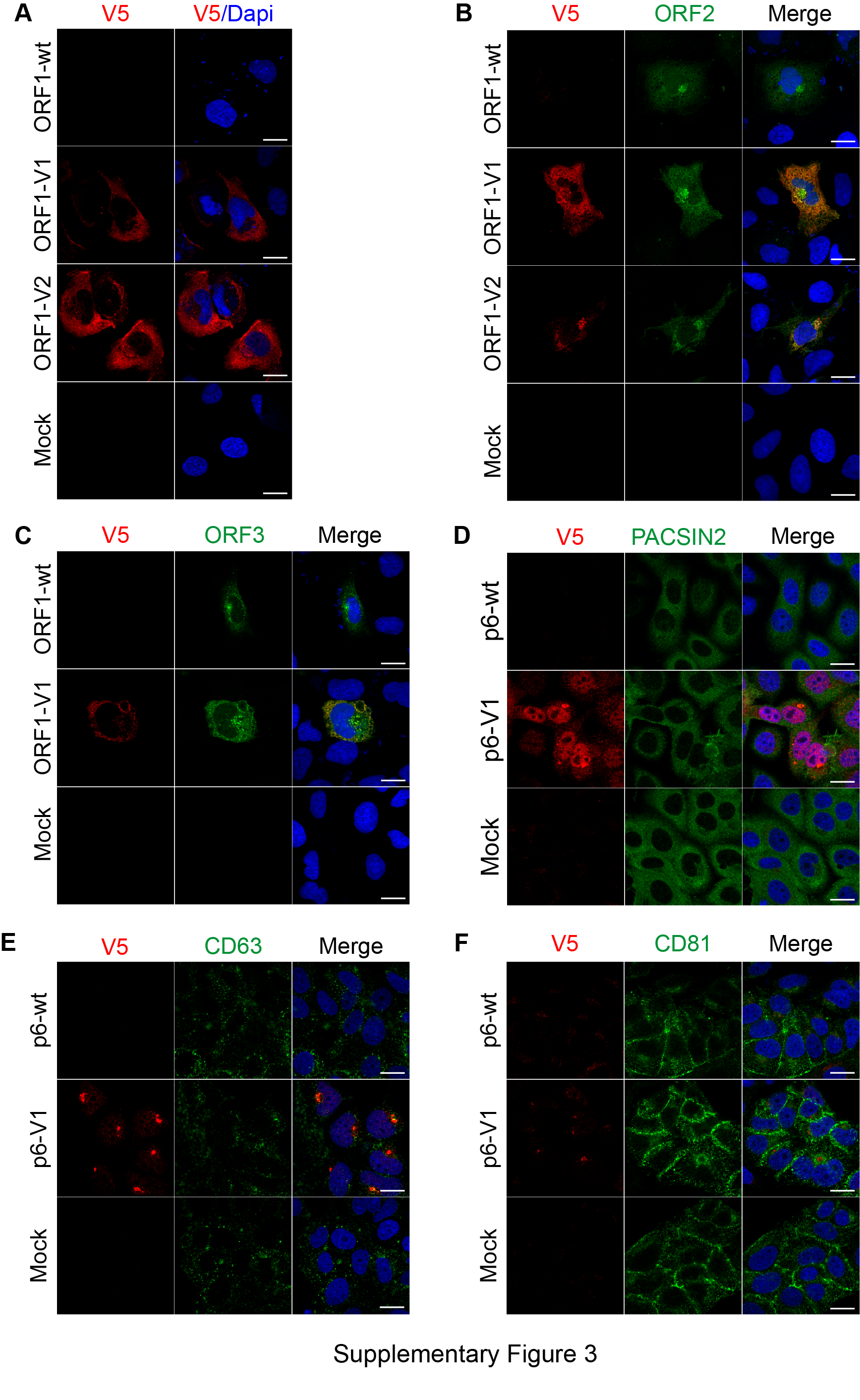
